# Multiplex editing of the *Nucleoredoxin1* tandem array in poplar: from small indels to translocations and complex inversions

**DOI:** 10.1101/2022.08.26.505498

**Authors:** Yen-Ho Chen, Shakuntala Sharma, William P. Bewg, Liang-Jiao Xue, Cole R. Gizelbach, Chung-Jui Tsai

## Abstract

The CRISPR-Cas9 system has been deployed for precision mutagenesis in an ever-growing number of species, including agricultural crops and forest trees. Its application to closely linked genes with extremely high sequence similarities has been less explored. Here, we used CRISPR-Cas9 to mutagenize a tandem array of seven *Nucleoredoxin1 (NRX1*) genes spanning ~100 kb in *Populus tremula* × *alba*. We demonstrated efficient multiplex editing with one single gRNA in 42 transgenic lines. The mutation profiles ranged from small indels and local deletions in individual genes to large genomic dropouts and rearrangements spanning tandem genes. We also detected complex rearrangements including translocations and inversions resulting from multiple cleavage and repair events. Target capture sequencing was instrumental for unbiased assessments of repair outcomes to reconstruct unusual mutant alleles. The work highlights the power of CRISPR-Cas9 for multiplex editing of tandemly duplicated genes to generate diverse mutants with structural and copy number variations to aid functional characterization.

## Introduction

The CRISPR-Cas9 is a powerful tool for genome engineering in a growing number of model and nonmodel species.^1,3^ CRISPR-Cas9 directed by short gRNAs (guide RNAs) targets genomic regions of interest to generate double-strand breaks (DSBs) for repair by the error-prone non-homologous end joining (NHEJ) pathway, although involvement of alternative repair pathways has also been reported.^4–8^ CRISPR-Cas9 has been extensively used for gene knockout (KO) or disruption of *cis*-regulatory elements in both plant and animal genomes.^9^ For long-lived perennial trees, efficient mutagenesis by CRISPR-Cas9 enables recovery of null mutants in the first generation for functional characterization with greater specificity and precision than previous gene silencing-based approaches.^4,10–12^

Despite its broad adoption, application of CRISPR-Cas9 to tandemly arrayed gene (TAG) editing has been limited. TAGs comprise ~10-20% of the gene content in plant and vertebrate genomes.^13–15^ Moreover, TAGs are enriched in functional categories associated with stress response and secondary metabolism, and they contribute to lineage-specific gene family expansion.^16–18^ Gene family expansion and subsequent homogenization occur because TAGs are often localized to chromosomal regions with high rates of recombination.^13,18^ However, the high degrees of coding and regulatory sequence similarity pose significant challenges to functional characterization of TAGs.^19^ Multigene mutations/lesions are necessary to address genetic redundancy, but screening for meiotic recombination between closely linked TAGs is difficult if not impossible even in model species.^19^

Genome engineering with zinc finger nucleases has been used to generate both small and large deletions as well as inversions in TAGs; however, the efficiencies across multiple TAG families were low.^20,21^ Efficient TAG editing by CRISPR-Cas9 has been reported in *Arabidopsis thaliana*^22,23^ and *Medicago truncatula*^24^ using two or more gRNAs to create multiple DSBs which can lead to large genomic dropouts in addition to short insertions and deletions (indels). Here we report efficient CRISPR-Cas9 mutagenesis of the *Nucleoredoxin1 (NRX1*) tandem array spanning ~100 kb in *Populus tremula* × *P. alba* INRA 717-1B4 (hereafter, 717 or hybrid poplar). We present evidence of multiple cleavages by a single gRNA in all 42 transgenic events, which resulted in diverse repair outcomes including small indels, large genomic dropouts, gene fusions, translocations, and inversions. We discuss analytical challenges associated with mutation profile determination at repetitive loci by different methods. The suite of mutants encompassing total nulls as well as those harboring in-frame fusion or wildtype alleles represent novel genetic materials for future functional characterization of NRX1 in hybrid poplar.

## Experimental Procedures

### CRISPR/Cas9 construct preparation and hybrid poplar transformation

A pair of oligos containing the gRNA sequence were used in conjunction with vector-specific primers (Table S1) for PCR amplification of *Medicago truncatula* U6 promoter and scaffold from the pUC-gRNA plasmid (Addgene #47024) using Q5® High-Fidelity 2X Master Mix (New England Biolabs). The two fragments were annealed and extended by overlapping PCR, digested with restriction enzymes *EcoRV* and *Eco*RI (New England biolabs), gel-purified with Zymoclean™ Gel DNA Recovery Kit (Zymo Research), and cloned into *SpeI* and *Swa*I digested p201N Cas9 plasmid (Addgene #59175). *E. coli* (DH5a) transformants were screened by colony PCR and positive clones confirmed by Sanger sequencing before being transformed into *Agrobacterium tumefaciens* strain C58/pMP90. Transformation of hybrid poplar 717 was performed as described by^25^ using WT and three previously produced transgenic lines, F10, F52, and F55 (also in 717 background)^26^. Regenerated plants were transplanted to soil, vegetatively propagated by single node cuttings and grown in a greenhouses as described.^27^

### PacBio long read assembly

The PacBio data of 717 was downloaded from NCBI SRA (PRJNA332358). A subset (SRR4228355-SRR4228446) of the dataset was used for assembly by Racon^28^ and Canu v1.4.^29^ All contigs were searched using *PotriNRX1* sequences to identify *NRX1*-containing contigs. The raw reads were mapped back to the contigs to check the assembly quality. Illumina resequencing reads of 717 (PRJNA266903) was mapped to the Chr10 long contig and visualized using Integrative Genomics Viewer ^30^ for manual curation.

### Mutation pattern determination by PCR and amplicon sequencing

Genomic DNA was extracted from leaf samples of tissue culture or greenhouse-grown plants as described.^31^ PCR was performed using either consensus primers that amplify all *NRX1* TAGs or genespecific primers (Table S1). Because of the sequence similarity, some primers are shared between subsets of TAGs and the *PtaNRX1.2* primer pair amplifies other TAGs with different sizes (see text). The specificity of the primer pairs was verified by Sanger sequencing of the corresponding PCR products from WT at Eurofins Genomics (Louisville, KY) or GENEWIZ/Azenta Genomics (South Plainfield, NJ). PCR products were analyzed by electrophoresis and visualized on a UV transilluminator.

Amplicon library was prepared as described ^25,32^ using multiple sets of primers. Amp-seq primer set 1 was used for the initial assessment and missed an SNP in *PtaNRx1.7*. Amp-seq primer set 2 was redesigned based on the curated 717 sequences for amplification of all *PtaNRX1s* except *PtaNRX1.2*. Individual plant samples were then barcoded in a second PCR and pooled equally into one library. A separate library was made using Amp-seq primer set 3 specifically for *PtaNRX1.2*. For the subset of greenhouse-grown mutant events, Amp-seq primer set 4 conserved in all seven TAGs was used. Amplicon libraries were sequenced on an Illumina MiSeq Nano flowcell (PE150) at the University of Georgia’s Georgia Genomics and Bioinformatics Core. Demultiplexed sequence data were analyzed by AGEseq.^33^ Initial analysis with only the native *NRX1* target sequences showed inconsistent results with many partial/chimeric edits, and manual curation suggested their origins from potential gene fusions. We therefore included all hypothetical gene fusions as potential targets for AGEseq analysis with 0% mismatch allowance due to the high degree of sequence similarity between the TAGs. The output was manually curated with noise thresholds determined based on WT patterns and sequencing depth of each sample. Where applicable, missing data were repeated by gene-specific PCR followed by sanger sequencing of purified PCR products.

### Target capture sequencing

High molecular weight genomic DNA was extracted from young leaves of greenhouse-grown plants using the CTAB method,^34^ and concentration determined by Qubit dsDNA HS assay kit (Invitrogen, Waltham, MA). For each of the samples, 700 ng genomic DNA in 55 μL ddH2O was sheared to ~550 bp using Covaris E220 (Woburn, MA) and size-selected using AMPure XP beads (Beckman Coulter, Indianapolis, IN). The fragmented genomic DNA was hybridized with xGen Lockdown probes (Table S2) tiled across *PtaNRX1.7* sequence (Integrated DNA Technologies or IDT, Coralville, IA), washed, and purified based on the manufacturer’s instructions before sample barcoding with xGen Stubby Adapter and Unique Dual Index Primers (IDT) and library preparation using KAPA HyperPrep Kit (Kapa Biosystems-Roche, Wilmington, MA). The library was then sequenced by Illumina MiSeq nano PE150 at the Georgia Genomics and Bioinformatics Core. The output sequences were then mapped to the *NRX1* tandem array and pseudogenes using BBmap implemented in Geneious Prime (San Diego, CA), with maximum indel size set at 100 kb, ambiguous mapping set at first best and a Kmer length of 15. *De novo* assembly was performed using Geneious assembler with high sensitivity and assembled contigs were examined by multiple sequence alignments with *PtaNRX1* sequences.

## Results

### CRISPR targeting and species diversity of the *NRX1* tandem array in poplars

The *Populus trichocarpa* Nisqually-1 reference genome v4.1 contains nine *NRX1* TAGs on chromosome 10 (chr10) (Fig. 1A). Eight of them, *PotriNRX1.1* to *PotriNRX1.8* (Potri.010G059401 to Potri.010G060200), represent putative full-length genes, while *PotriNRX1.9p* (Potri.010G060300) is 5’-truncated and likely a pseudogene. These NRX1 TAGs share high levels of nucleotide coding sequence identity (Fig. S1A). The *P. trichocarpa* v4.1 genome contains another pseudogene on the homoeologous chr8, *PotriNRX1.10p* (Potri.008G175600), which is both 5’- and 3’-truncated. We followed our variant-free gRNA design principles^35,36^ and identified a spacer sequence within the second exon that is predicted to target all *NRX1* TAGs on chr10 (Fig. 1B).

**Figure 1.**
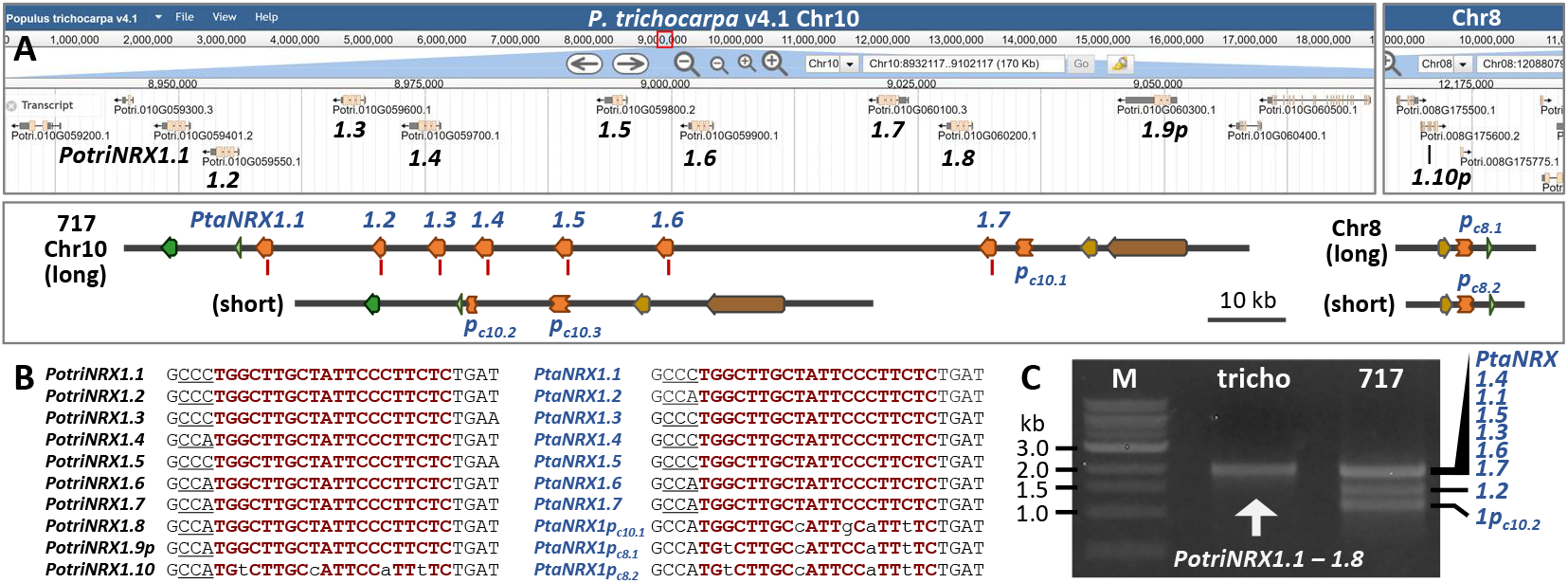
Species diversity of the *NRX1* tandem array **A.** A genome browser view of the *P. trichocarpa* (v4.1) *PotriNRX1* tandem array on chromosome 10 (chr10, top left) and a pseudogene on homoeologous chr8 (top right). The corresponding genomic regions of *P. tremula* x *alba* INRA 717-1B4 (717) are shown in the bottom panel, with the two homologous chromosomes depicted based on PacBio (long and short) contigs. *NRX1* genes are shown in orange arrows and pseudogenes (p) in truncated boxes. Red lines denote gRNA target sites. Flanking genes are color-coded for collinearity. **B.** Alignments of the gRNA target sites (red) based on *P. trichocarpa* (left) and 717 (right) sequences. The protospacer adjacent motif (PAM) is underlined. **C**. Distinct banding patterns of multi-gene PCR products between the two genotypes using consensus *NRX1* primers. M, molecular weight ladders, tricho, *P. trichocarpa*.

*Agrobacterium tumefaciens-mediated* transformation was performed in four 717 backgrounds, wild type (WT) and three SA hyper-accumulating lines (F10, F55, and F52) from a previous study.^26^ We obtained 42 independent events: 14 in WT (group A), 6 in F10 (group B), 12 in F52 (group C), and 10 in F55 (group D). We designed consensus primers flanking the gRNA target site of all TAGs for short amplicon sequencing using tissue cultured plants. Initial data analysis revealed poorer-than-expected editing efficiency across the TAGs because CRISPR-Cas9 mutagenesis rates in the hybrid poplar system typically approach 100%.^25,32,36,37^ To probe this further, we designed another pair of consensus primers for long PCR spanning 5’-UTR to the fourth exon (~2 kb) of the TAGs. We observed unexpected banding pattern variations between the *P. trichocarpa* Nisqually-1 reference and 717 hybrid poplar samples (see below). Specifically, the 717 WT yielded three bands approximately 2 kb, 1.5 kb, and 1 kb, whereas the *P. trichocarpa* Nisqually-1 sample produced the expected ~2 kb band cluster (Fig. 1C). This suggested greater than expected species diversity in the *NRX* TAGs, which might have confounded amplicon sequencing data analysis.

To investigate the species diversity, we mined PacBio data from the 717 genome sequencing project by the U.S. Department of Energy Joint Genome Institute (NCBI SRA BioProject PRJNA332358). We identified two contigs that span the *NRX1* tandem array (Data S1), one considerably longer (~259 kb, 217 reads) than the other (~105 kb, 175 reads). We used short Illumina reads from 717 resequencing data^35^ for error correction to generate 717-specific *NRX1* sequences (Fig. S2). We identified seven *NRX1* TAGs (*PtaNRX1.1* to *PtaNRX1.7*) along with a pseudogene on the long contig, and only *NRX1* remnants on the short contig (Fig. 1A). Based on homology and collinearity of flanking genes, we concluded that the two contigs represent homologous chr10s with distorted *NRX1* representation. We also identified two PacBio contigs that harbor truncated *NRX1* sequences homologous to the Chr8 pseudogene *PotriNRX1.10p* (Data S1). These predictions were confirmed in the recently released, haplotype-resolved 717 genomes (HAP1 and HAP2 v5.1) in Phytozome v13 (Table S3). Hereafter, we refer to the 717 *NRX1 <u>p</u>*seudogenes (and their alleles) on *<u>c</u>*hr*<u>10</u>* as *PtaNRX1p_c10.1_* (long contig), *PtaNRX1p_c10.2_* and *PtaNRX1p_c10.3_* (short contig), and on chr8 as *PtaNRX1p_c81_* and *PtaNRX1p_c82_*. As expected, the seven *PtaNRX1* TAGs share high levels of coding sequence identities (Fig. S1B). Despite the unexpected copy number variation, our SNP-aware gRNA design based on the *P. trichocarpa* genome matched a consensus region of the seven 717 *NRX1* TAGs but not the pseudogenes (Fig. 1B). We concluded that this gRNA should be effective for CRISPR-Cas9 mutagenesis in 717 and that the poor editing results from the initial analysis were likely caused by TAG variation between species.

We re-assessed the PCR results mentioned above using the curated 717 *NRX1* sequences. Of note is *PtaNRX1.2* which harbors an in-frame deletion downstream of the gRNA target site (Fig. S2) and is predicted to yield a 1,480 bp product with the long PCR primers. This primer pair can also amplify pseudogene *PtaNRX1p_c10.2_* with a product of ~1 kb. The other TAGs are expected to yield PCR products of ~2 kb (Fig. S2, hereafter, referred to as multi-gene PCR). These sequence-based predictions explain the multi-gene PCR banding pattern discrepancies between 717 and *P. trichocarpa* (Fig. 1C).

### Mutation pattern determination by PCR and amplicon deep sequencing

Using the curated 717 *NRX1* sequences, we designed two new sets of primers for amplicon sequencing (Amp-seq primer sets 2 and 3, Table S2). Set 2 primers are conserved among six *PtaNRX1* TAGs (all but the internal-truncated *PtaNRX1.2*) and pseudogene *PtaNRX1p_c8_*(both alleles), while Set 3 primers favor *PtaNRX1.2* amplification under our PCR conditions (161 bp versus >700 bp for other TAGs). Because of the challenge associated with short read mapping among highly similar sequences, and because of the potential for intramolecular fusion between two CRISPR-Cas9-cleavage sites (see below), we included all hypothetical fusions between *PtaNRX* TAGs as potential targets for amplicon data analysis by AGEseq^33^ (see Methods). Across the 294 potential target sites in all four transgenic groups (7 sites × 42 lines), native (unedited) alleles were detected only at seven sites (2%, green color in Figs. 2A and 2B, Data S2), suggesting highly efficient editing of the *PtaNRX1* tandem repeats. Approximately 32% of the target sites harbor small insertions and deletions (indels, orange color in Figs. 2A and 2B), with single-base deletions (−1) and insertions (+1) being most common (Fig. 2B).

**Figure 2.**
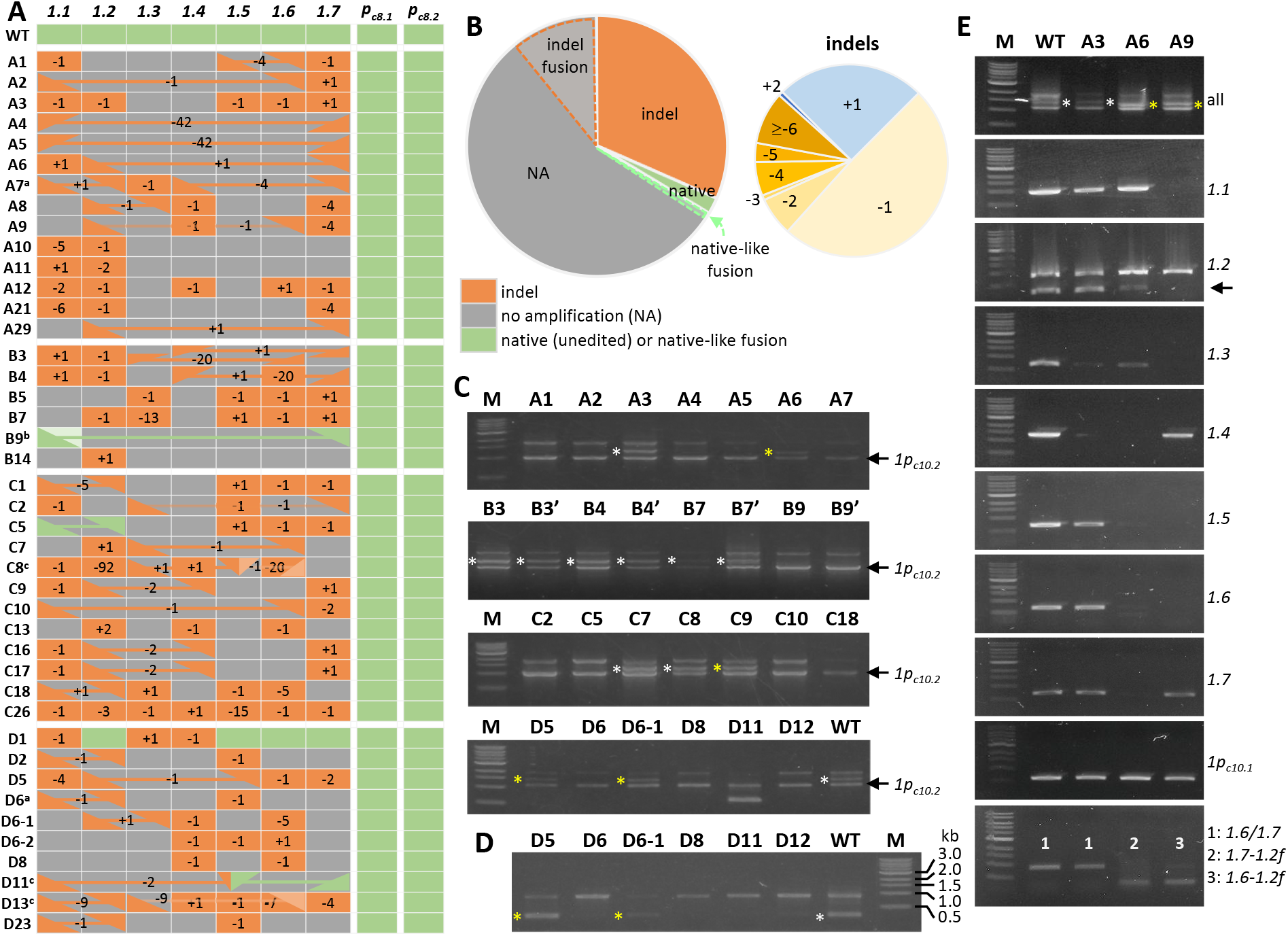
Mutation profiles of the *NRX1* tandem repeats **A.** Mutation patterns as determined by amplicon sequencing. The seven *NRX1* tandem genes on Chr10 and the two pseudogene alleles on Chr8 are shown in columns with plant lines in rows. Mutation types are color-coded, with indel patterns noted. Linked triangles represent fusion alleles, with the fusion orientation denoted as from top of cell (5’, 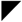 or 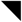) to bottom of cell (3’, 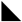 or 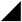). Curated edits are as follows: ^a^ *PtaNRX1.2-like* reads were detected in A7 and D6, but not by multi-gene or gene-specific PCR. ^b^ The *PtaNRX1.1*-like allele (light green) in B9 was confirmed to be a fusion between *PtaNRX1.7* and *PtaNRX1.1*. ^c^ Different shades of orange (C8 and D13) or two triangles in a cell (C8, D11) denote the involvement of some alleles in complex rearrangements. **B.** Pie chart depiction of the overall editing patterns based on amplicon sequencing (left). The slices with dashed outlines represent the portion of indel or native/native-like fusion alleles spanning NA alleles. The breakdown of indel types from both native and fusion alleles is shown on the right. **C.** Multi-gene PCR banding patterns of representative plants. The truncated *PtaNRX1.2* is readily identifiable as the middle band (white asterisks for native alleles, yellow asterisks for fusion alleles with *3’-PtaNRx1.2*). The lower band is from the non-target pseudogene *PtaNRX1p_c10.2_* and serves as loading control. The group B panel includes duplicate samples from independent plants (B3’, B4’, B7’ and B9’). **D.** Gene-specific PCR for *PtaNRX1.2* (344 bp) corroborated the multi-PCR results shown in C. Native or fusion *PtaNRX1.2* alleles are marked with white and yellow asterisks, respectively. This prime pair also amplified *PtaNRX1.1/1.3/1.4/1.5* of ~900 bp. **E.** PCR analysis of WT, A3, A6 and A9 using consensus or gene-specific primers. The bottom panel was performed using a hybrid (*PtaNRx1.6/1.7* forward and *PtaNRX1.2* reverse) primer pair intended to amplify *PtaNRX1.6-1.2* (A9) and *PtaNRX1.7-1.2* (A6) fusions, at ~350 bp. The reverse primer has two mismatches for *PtaNRX1.6/1.7* and produced non-specific products of ~910 bp.

Many of the target sites (66%) yielded no amplification (NA, grey color in Figs. 2A and 2B), likely due to the absence of primer-binding site(s). This assessment was strengthened by amplification in all plant samples of the two non-target *PtaNRX1p_c8_* alleles, which served as a built-in quality control for PCR (*i.e.*, NA was not due to PCR failure). NA alleles were found in all but two transgenic lines, and often spanned two or more consecutive TAGs (Fig. 2A). Such patterns may be indicative of large genomic dropouts resulting from simultaneous Cas9 cleavage at two or more adjacent target sites, as reported previously.^25^ Indeed, multi-gene (long) PCR with consensus primers showed that the intensity of the ~2 kb band cluster was inversely correlated with the number of NA alleles using the non-target *PtaNRX1p_c10.2_* band as loading control (Figs. 2A and 2C). Lines A4, A5, and B9 represent the extremes, each exhibiting a faint ~2 kb band and 5-6 consecutive NA alleles. The loss of *PtaNRX1.2* in the surveyed transgenic events was readily discernable (Fig. 2C, asterisks). The results were corroborated by another set of primers that amplified a 344 bp *PtaNRX1.2* fragment (in addition to ~900 bp from *PtaNRX1.1/1.3/1.4/1.5*) (Fig. 2D). The only exceptions were A7 and D6 where *PtaNRX1.2-like* sequences were detected by amplicon sequencing (Data S2) even though both lines were negative for *PtaNRX1.2* in multi-gene and gene-specific PCR (Fig. 2C). The discrepancy suggested other genomic rearrangements following CRISPR-Cas9 editing. Unusual PCR banding patterns, such as those detected in D11 (Fig. 2C), are also indicative of large deletion(s).

### Frequent detection of *NRX1* fusion alleles between different cleavage sites

During manual examination of amplicon sequencing results, we noticed multiple cases of ambiguously mapped reads representing hybrid sequences between distinct *NRX1* genes. This prompted us to include hypothetical target sequences between all possible fusions of sequential (adjacent or non-adjacent) TAGs for AGEseq analysis. We identified 34 fusion alleles (denoted as linked triangles, Fig. 2A) in >60% of the transgenic population. The number is likely an underestimate as only those with amplicon primer binding sites were captured in this analysis. The majority of the fusion alleles harbor frameshift indels, though in-frame fusion events resulting in native-like alleles were detected in several instances (*e.g.*, C5 and D11). More than 60% of the fusion alleles involved *PtaNRx1.1* and/or *PtaNRX1.2*, with the *PtaNRX1.2-1.1* fusion being most common (Fig. 2A). Fusions between non-adjacent TAGs frequently coincided with loss of intervening copies (NA alleles). As an example, fusion between *5’-PtaNRX1.7* and *3’-PtaNRX1.2* in A6 resulted in the loss of *PtaNRX1.3* through *PtaNRX1.6*, with an estimated dropout of ~77 kb. Accordingly, A6 was PCR-positive for only *NRX1.1* and the non-target control *PtaNRX1p_c10_._2_* (Fig. 2E). The *PtaNRX1.7-1.2* fusion yielded a fragment that is ~100 bp shorter than *PtaNRX1.2* by multi-gene PCR (Figs. 2C and 2E, yellow asterisks). This fusion was also detected by a hybrid primer pair matching 5’-*PtaNRX1.6/1.7* and *3’-PtaNRX1.2* (Fig. 2E, bottom panel).

### Long-term stability of CRISPR-Cas9 edits

A subset of 12 transgenic lines from the four genetic backgrounds (three each) and WT plants were transplanted to soil and grown in a greenhouse. Plants were further propagated by cuttings and maintained for two years before DNA analysis to assess the stability of CRISPR-mediated *NRX1* TAG editing outcomes. We performed amplicon sequencing using new primers (Amp-seq primer set 4) that are consensus for all seven TAGs, as well as multi-gene and gene-specific PCR. We found the same mutation patterns as in cognate samples from tissue culture (Fig. 3B, Data S2 and S3), including inframe fusion alleles with intact gRNA target sequence (*i.e.*, no further editing was observed). CRISPR editing outcomes were therefore stable through multiple rounds of vegetative propagation as previously observed.^4^ Despite the clonal stability, conflicting results between amplicon sequencing and genespecific PCR were noted in both rounds. For instance, B14 showed six NA alleles by amplicon sequencing, but only three by gene-specific PCR. Such discrepancies could be caused by mutations impacting primer-binding sites or other complex rearrangements and reflect limitations of PCR-based assays.

**Figure 3.**
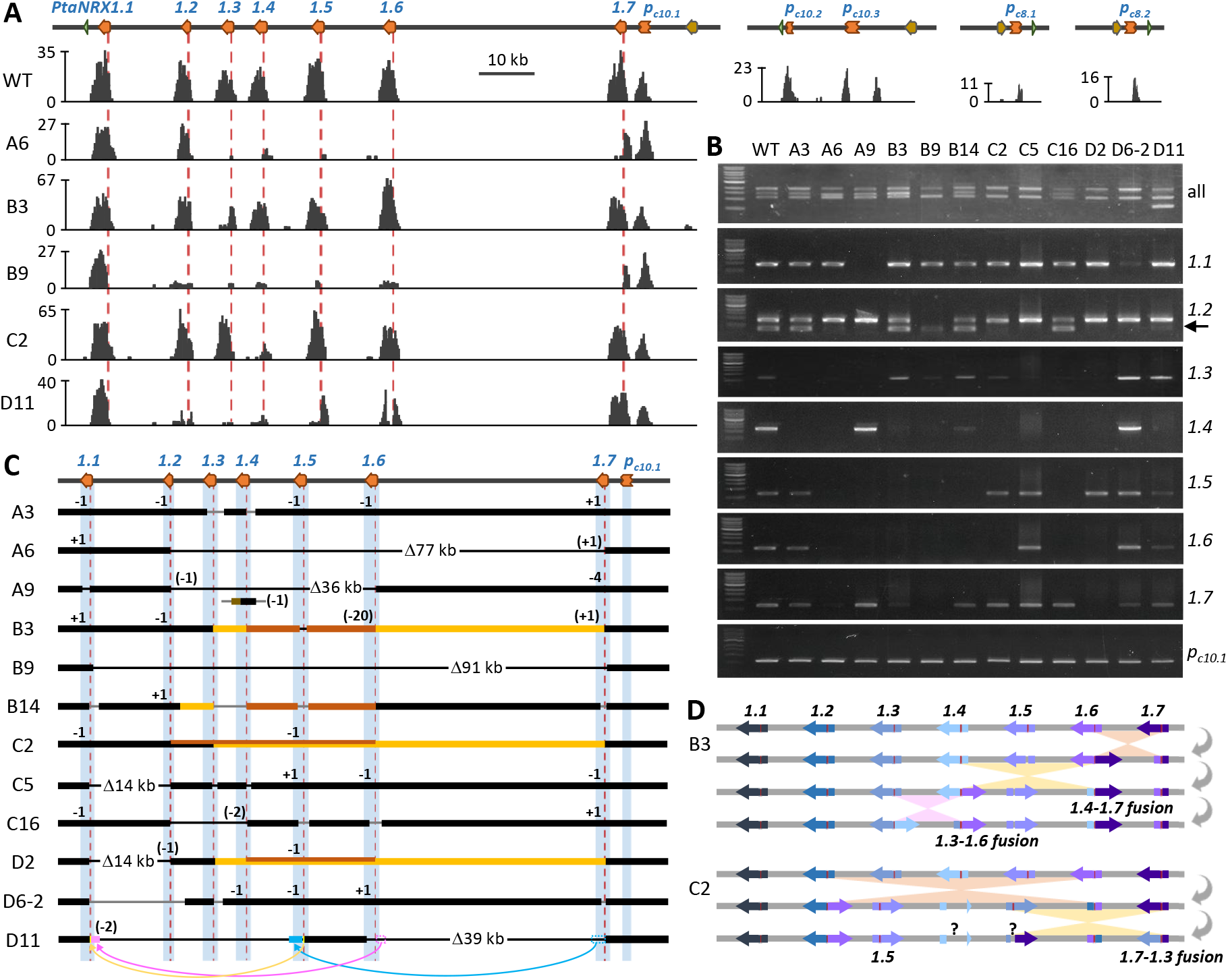
Target capture sequencing and PCR analysis of greenhouse plants **A.** Sequence coverage profiles across the *NRX1* tandem array on Chr10. The genomic region is shown on top and representative read mapping profiles are shown below (Y axis, read counts). The read mapping profiles to the *NRX1* pseudogenes (homologous Chr10 and two homologous Chr8 alleles) are also shown for the WT sample. Red dashed lines denote the gRNA target site. **B.** PCR analysis of greenhouse-grown WT and 12 mutants using consensus or gene-specific primers. **C.** Summary of the editing outcomes for the panel of 12 mutants based on amplicon sequencing, PCR and capture sequencing. Blue columns and read dashed lines denote the NRX1 gene space and the gRNA target site, respectively. Black bars represent the tandem array region with back and gray lines connecting confirmed or unknown fusion conjunctions, respectively. Yellow and orange bars in B3, B14, C2 and D2 denote distinct inversion events. The lower track for A9 shows a translocated fusion allele between *5’-PtaNRX1.4* (black bar) and pseudogene 3’*-PtaNRX1p_c10.3_* (brown bar), presumably on the alternative chr10. Suspected rearrangement events in D11 are shown by colored (open or solid) bars and arrows. **D.** Schematic illustration of sequential inversion events that resulted in the flipped fusion alleles in B3 and a correctly oriented fusion allele in C2. Question marks denote as-yet-undetermined junctions.

### Target capture sequencing revealed multiple cases of complex rearrangements

We next carried out target capture sequencing to enrich the *NRX1* locus (see Methods) in order to probe into some of the unusual or as-yet-unexplainable mutation patterns. We first validated the capture sequencing workflow with the WT sample and demonstrated successful enrichment of *NRX1* TAGs and, to a lesser extent, *NRX1* pseudogenes (Fig. 3A). Analysis with mutant samples confirmed many of the NA alleles previously called by amplicon sequencing (Figs. 2A and 3A). For instance, line A6 showed a dearth of mapped reads for *PtaNRX1.3* through *PtaNRX1.6* (Fig. 3A), which is consistent with a large genomic dropout between *PtaNRX1.2* and *PtaNRX1.7* (Figs. 3B and 3C). Line B9 also harbors a large genomic dropout between *PtaNRX1.1* and *PtaNRX1.7* (~91 kb), supported by a scarcity of reads for the intervening TAGs (Fig. 3A). We further confirmed many of the amplicon sequencing-identified fusion alleles with paired reads spanning the fusion junctions (denoted by black lines, Fig. 3C). An example is the *PtaNRX1.7-1.2* fusion in line A6 (Fig. 3C), which was independently confirmed by PCR (Fig. 2E) and Sanger sequencing (Data S4). However, several fusion junctions remain undefined.

We then explored *de novo* assembly of *NRX1*-enriched sequence reads for unbiased assessments of editing outcomes that escaped PCR-based detection. Using B9 as a test case, we recovered a previously undetected fusion junction between 5’*-PtaNRX1.7* and *3’-PtaNRX1.1* (Data S4). This in-frame fusion occurred ~120 bp upstream of the gRNA target site or ~60 bp upstream of the amplicon forward primer sequence, thus explaining its (mis)identification as unedited *NRX1.1* by amplicon sequencing (Fig. 2A). In the case of A9, *de novo* assembly revealed the −20 fusion allele as *PtaNRX1.6-1.2*, as opposed to *PtaNRX1.3-1.2* called by amplicon sequencing based on a single mismatch at the PAM site, which might have been introduced during fusion/repair. This fusion was independently confirmed by PCR (Fig. 2E) and Sanger sequencing (Data S4). *De novo* assembly indicated another fusion between a mutated *PtaNRX1.4* (likely derived from the genomic drop out of the *PtaNRX1.6-1.2* fusion event) and pseudogene *PtaNRX1p_c10.3_* on the alternative chr10 (Fig. 3C, Data S4). This implied a new genomic context of the edited *PtaNRX1.4* via a translocation event in A9.

In another example, D11 was predicted by amplicon sequencing to harbor two fusion alleles as the only remaining TAGs, both involving *PtaNRX1.5; PtaNRX1.5-1.1* (−2) and *PtaNRX1.5-1.7* (in-frame fusion). However, capture sequencing recovered a high number of reads mapping to *PtaNRX1.7* (Fig. 3A), as well as paired reads spanning different genes (5’-3’: *PtaNRX1.6-PtaNRX1.1, PtaNRX1.6-PtaNRX1.5, PtaNRX1.7-PtaNRX1.6*, in addition to *PtaNRX1.5-PtaNRX1.7*). Furthermore, D11 yielded a truncated fragment (~600 bp in length) by multi-gene PCR (Figs. 2C and 3B) and an unexplainable PCR product by primers designed for *PtaNRX1.3* (Fig. 3B). *De novo* assembly of capture sequencing data from D11 recovered a 3-way fusion allele consisting of *5’-PtaNRX1.6*, a short *PtaNRX1.5* fragment (~80 bp) immediately upstream of the gRNA target site, and *3’-PtaNRX1.1* downstream of the gRNA target (Fig. 3C, Data S4). This explained its (mis)identification as a *PtaNRX1.5-1.1* fusion by amplicon sequencing as delineated by primers (Fig. 2A, Data S2 and S3). *De novo* assembly confirmed the in-frame fusion allele *PtaNRX1.5-1.7* in a flipped orientation with respect to the genomic configuration (*i.e*., 5’*-PtaNRX1.5* and 3’*-PtaNRX1.7*, Fig. 3C). This fusion resulted in unexpected primer-binding sites shared with *PtaNRX1.3* (Table S2) and explains the mysterious amplification noted above (Fig. 3B). Sanger sequencing of this PCR product validated its identity as a *PtaNRX1.5-1.7* fusion (Data S4). Finally, *de novo* assembly identified a third fusion allele consisting of *5’-PtaNRX1.7* and *3’-PtaNRX1.6*, with a ~1.3 kb deletion downstream of the gRNA cleavage site. This resulted in loss of the reverse primer binding site for amplicon sequencing, rendering the fusion allele undetected. The internal deletion also explains the unusually small fragment (~600 bp) from multi-gene PCR (Figs. 2C and 3B), which was confirmed by Sanger sequencing as *PtaNRX1.7-1.6* fusion (Data S4). We note the unusual configurations of these fusion alleles with respect to the order and orientation of the cognate TAGs; their genomic contexts remain unknown at the present time.

In several mutant lines, we observed unusual mapping orientation and/or distance between paired reads, suggesting potential inversion. An example is B14 which harbors at least two inversions (Fig. 3C), one between (upstream of) *PaNRX1.2* and *PaNRX1.3* and the other between *PtaNRX1.4* and *PtaNRX1.6*, in addition to the loss of *PtaNRX1.5* in the latter. This likely underlies the discrepancy between amplicon sequencing, gene-specific PCR and capture sequencing read distribution, as well as other ambiguously mapped reads in this line (Figs. 2A, 3B, and 3C, Data S2 and S3). B3 harbors two flipped fusion alleles which could have arisen from three sequential inversion events as illustrated in Fig. 3D. The *PtaNRX1.4-1.7* fusion was unexpectedly amplified by primers intended for *PtaNRX1.3* (Fig. 3B), and was verified by Sanger sequencing (Data S4). We detected two putative double inversion events in C2 and D2, with one inversion nested within another (denoted by yellow and orange lines, Fig. 3C). In the case of C2, the rearrangements resulted in nonproductive gene fusions at the junctions (opposing orientation of the two joined fragments), except for the *PtaNRX1.7-1.3* fusion which retained the proper orientation with a typical indel (−1) as identified by amplicon sequencing. This allowed us to reconstruct the sequence of events that led to the double inversion in C2 (Fig. 3D): first inversion between *PtaNRX1.2* and *PtaNRX1.6* followed by a second inversion between *PtaNRX1.3* and *PtaNRX1.7* effectively restored the proper orientation for the *5’-PtaNRX1.7* and *3’-PtaNRX1.3* fusion to permit detection by amplicon-sequencing.

## Discussion

Here we report efficient multiplex editing of tandemly duplicated and highly homologous *NRX1* genes using a single gRNA. All 42 transgenic lines we characterized harbor mutations in the tandem array. The editing outcomes are diverse, ranging from small indels and local deletions impacting individual TAGs, to large genomic dropouts and complex rearrangements spanning multiple TAGs. Across all target sites, NA alleles were the most common mutation type, twice as frequent as small indels based on amplicon sequencing. This contrasts with single-target editing where small indels predominate,^38,39^ but is consistent with the recent report of a positive correlation between NA allele frequency and tandem gene (target) copy number.^25^ At the individual plant level, NA alleles were found in nearly all transgenic lines and frequently spanned consecutive TAGs. In many cases, consecutive NA alleles are associated with gene loss (genomic dropout) due to simultaneous Cas9 cleavages at two or more target sites, as exemplified by A6 and B9 (Figs. 3A and 3C). Repair of two or more DSBs can lead to gene fusions, which were detected in two-thirds of the mutant population. While fusions of adjacent TAGs were common (12 of 34 detected fusion alleles), the same number of fusions occurred between well separated (>35 kb), non-consecutive TAGs (Figs. 2A and 3C).

Capture sequencing revealed several cases of gene inversions and complex rearrangements that escaped PCR-based detection. The high degree of sequence similarity among *NRX1* TAGs (and their pseudogenes) means homologous templates were readily available to facilitate homology-directed repair (HDR) or microhomology-mediated end joining (MMEJ) of Cas9-induced DSBs^8,40,41^. *De novo* assembly of target-enriched sequencing reads allowed us to reconstruct several repair junctions and provided evidence for HDR involvement in some of the repair outcomes. A notable example is A9 where the mutated *PtaNRX1.4* from the dropout of the *PtaNRX1.6-1.2* fusion was translocated and fused with the pseudogene *PtaNRX1p_c10.3_* on alternative chr10, with extensive sequence homology at one junction. This is reminiscent of CRISPR-Cas9-mediated chromosomal reshuffling in yeast by targeting repetitive LTRs (long terminal repeats) to induce multiple DSBs and subsequent translocations by HDR using (uncut) LTR as template.^40^ Also unrelated to TAG editing, CRISPR-based chromosome engineering has been used to generate large-scale deletions, inversions, and/or translocations in yeast, plant, and animal systems, including structural rearrangements commonly detected in cancer cells.^8,24,41–46^ Similarly, CRISPR editing of gene clusters with multiple distinct gRNAs has resulted in DNA fragment deletions and inversions in mice and human cells.^8^ Taken together, the present work and our recent study^25^ both showed that a single gRNA is effective in multiplex editing of tandem duplicates, with mutation outcomes similar to those achieved by multiple gRNAs. The risk of off-target editing is also lowered when a single gRNA can be used to generate multiple DSBs.

Our work highlights both the efficiency and complexity of multiplex editing at the *NRX1* TAGs. The combination of high copy number and high sequence homology of TAGs presents multiple technical challenges for mutation mapping. An assortment of methods was used to characterize the mutants, each with its own advantages and limitations. Multi-gene and gene-specific PCR can detect large structural variations (*e.g.*, altered banding patterns in D11) but has limited resolution unless coupled with cloning and sequencing. Gene-specific primers can be difficult to find among nearly identical TAGs, and we had to rely on different combinations of consensus primers and/or amplicon lengths to differentiate some TAG members. A drawback is their unknown specificity for (unexpected) fusion alleles. For instance, a forward primer conserved for *PtaNRX1.1/1.2/1.3/1.4/1.5* and a reverse primer for *PtaNRX1.3/1.7* amplified not only *PtaNRX1.3* but also *PtaNRX1.5-1.7 (e.g.*, D11), *PtaNRX1.5-1.3 (e.g.*, C8), *PtaNRX1.4-1.7 (e.g.*, B3 and B4), and *PtaNRX1.1-1.7 (e.g.*, A4 and A5) fusion alleles, and the true identity of each can only be ascertained by sequencing.

Amplicon sequencing is powerful for multi-allele detection when consensus primers are used. The limitation lies in the short sequencing length (Illumina PE150 in this case) which can restrict discrimination between highly similar TAGs (*e.g*., some native and fusion alleles are identical or differ by only one or two SNPs). Increasing sequencing lengths can improve resolution, but costs increase as well. Other limitations of PCR-based assays still apply, as edits outside of the amplicon will be missed (*e.g*., *PtaNRX1.7-1.1* fusion in B9). Additional challenges associated with amplicon deep sequencing include underappreciated complexity of PCR primer specificity, such as variable effects of a single mismatch at the 3’ end as reported previously.^47^ This gives rise to data noise that warrants manual curation. Finally, capture sequencing allows target enrichment across the tandem array (and their pseudogenes) for unbiased assessment of mutation profiles. This enabled discovery of unusual fusion, inversion, and translocation events. While short read mapping to repetitive regions such as tandem repeats remains a technical challenge,^48^ increasing sequencing length and depth should alleviate some of the issues. The multipronged approach can also ensure absence of other “hidden” translocations at unexpected loci. For instance, the dearth of mapped reads in regions flanked by fused genes (*e.g*., A6 and B9, Fig. 3A) provides evidence that the intervening fragments were lost, whereas high read counts at NA alleles called by amplicon sequencing may signify other structural rearrangements. Finally, long read sequencing technologies with improved base calling accuracy hold promise to expedite mutation profile determination at repetitive loci.

In conclusion, we demonstrated the effectiveness of a single consensus gRNA for multiplex editing of TAGs spanning ~100 kb in the hybrid poplar. Of the mutant population we generated, at least 31 transgenic lines (5-10 per parent group) are predicted to be null mutants for *NRX1*. The three in-frame, native-like fusion alleles (*PtaNRX1.7-1.1* in B9, *PtaNRX1.2-1.1* in C5 and *PtaNRX1.5-1.7* in D11) represent novel *NRX1* genes owing to recombination of coding sequences as well as upstream, downstream and intronic sequences known to harbor *cis* regulatory elements that modulate gene expression.^49^ Tandemly duplicated genes have long-established roles in taxon-specific stress adaptation and they are known to undergo homogenization that contributes to functional diversification.^13,17^ Results from this study suggest that different CRISPR-based/Cas9 editing strategies can be designed to generate a wide spectrum of mutants with structural or copy number variations to aid functional investigation of TAGs.

## Author Contributions

C.-J.T conceived the project and designed the CRISPR experiment, S.S. generated transgenic plants and performed initial analysis, L.-J.X. performed PacBio assembly, Y.-H.C. performed all other mutant characterization and analyzed data with assistance from W.P.B and C.R.G on amplicon sequencing, Y.-H.C. and C.-J.T. wrote the manuscript with contribution from S.S.

## Acknowledgements

The authors thank Gilles Pilate of the Institut National de la Recherche Agronomique, France for providing hybrid poplar clone INRA 717-1B4, Wayne Parrott for p201N-Cas9 and pUC-gRNA plasmids, Xi Gu for an earlier attempt at *de novo* assembly of *PtaNRX1* cDNAs, and the Georgia Genomics and Bioinformatics Core for Illumina NextSeq sequencing. The 717 PacBio data and the haplotype genomes were generated by the US Department of Energy Joint Genome Institute, which is a DOE Office of Science User Facility supported by the Office of Science of the US Department of Energy under contract no. DE-AC02-05CH11231. The authors declare no conflicts of interest.

## Supporting Information

**Table S1.** Primers used in this study.

**Table S2.** xGen lockdown probes used in *NRX1*-capture sequencing.

**Table S3.** *PtaNRX1* gene models in the two 717 haplotype genomes.

**Figure S1.** Coding sequence identity matrix among poplar *NRX1* TAGs.

**Figure S2.** Multiple sequence alignments of *PtaNRX1* genes.

**Data S1.** *NRX1*-containing PacBio contigs of chr10 and chr8.

**Data S2.** Mutation patterns determined by amplicon sequencing of 42 mutant lines from tissue culture.

**Data S3.** Mutation patterns determined by amplicon sequencing of 12 mutant lines from greenhouse.

**Data S4.** Representative fusion allele sequences.

## Notes

**Funding**, The work was funded in part by Department of Agriculture, National Institute of Food and Agriculture (grant no. 2015-67013-22812), the Center for Bioenergy Innovation, a US Department of Energy Research Center supported by the Office of Biological and Environmental Research in the DOE Office of Science. Y.-H.C. received an Innovative and Interdisciplinary Research Grant from the Graduate School and a Palfrey Student Research Grant from the Department of Plant Biology, University of Georgia. C.R.G. was partly supported by National Science Foundation (REU Site grant no. DBI-1946937).

### Competing Interest Statement

The authors have declared no competing interest.

### Summary of Updates

minor text revision to include references on complex genomic rearrangements commonly detected in cancers

## References

1. Puchta H, Jiang J, Wang K, et al. Updates on gene editing and its applications. Plant Physiology 2022;188(4):1725–1730, doi:10.1093/plphys/kiac032

2. Brandt K, Barrangou R. Applications of CRISPR technologies across the food supply chain. Annual Review of Food Science and Technology 2019;10(1):133–150, doi:10.1146/annurev-food-032818-121204

3. Wang H, Russa ML, Qi LS. CRISPR/Cas9 in genome editing and beyond. Annual Review of Biochemistry 2016;85(1):227–264, doi:10.1146/annurev-biochem-060815-014607

4. Bewg WP, Ci D, Tsai C-J. Genome editing in trees: From multiple repair pathways to long-term stability. Frontiers in Plant Science 2018;9(1732, doi:10.3389/fpls.2018.01732

5. Gehrke F, Schindele A, Puchta H. Nonhomologous end joining as key to CRISPR/Cas-mediated plant chromosome engineering. Plant Physiology 2022;188(4):1769–1779, doi:10.1093/plphys/kiab572

6. Lemos BR, Kaplan AC, Bae JE, et al. CRISPR/Cas9 cleavages in budding yeast reveal templated insertions and strand-specific insertion/deletion profiles. Proceedings of the National Academy of Sciences 2018;115(9):E2040–E2047, doi:10.1073/pnas.1716855115

7. van Overbeek M, Capurso D, Carter Matthew M, et al. DNA repair profiling reveals nonrandom outcomes at Cas9-mediated breaks. Molecular Cell 2016;63(4):633–646, doi:10.1016/j.molcel.2016.06.037

8. Li J, Shou J, Guo Y, et al. Efficient inversions and duplications of mammalian regulatory DNA elements and gene clusters by CRISPR/Cas9. J Mol Cell Biol 2015;7(4):284–98, doi:10.1093/jmcb/mjv016

9. Wang JY, Doudna JA. CRISPR technology: A decade of genome editing is only the beginning. Science 2023;379(6629):eadd8643, doi:doi:10.1126/science.add8643

10. Tsai C-J. Genome editing of woody perennial trees. In: Genome Editing For Precision Crop Breeding. (Willmann MR. ed.) Burleigh Dodds Science Publishing: London; 2021.

11. Wang X, Tu M, Li Z, et al. Current progress and future prospects for the clustered regularly interspaced short palindromic repeats (CRISPR) genome editing technology in fruit tree breeding. Critical Reviews in Plant Sciences 2018;37(4):233–258, doi:10.1080/07352689.2018.1517457

12. Goralogia GS, Redick TP, Strauss SH. Gene editing in tree and clonal crops: progress and challenges. In Vitro Cellular & Developmental Biology - Plant 2021;57(4):683–699, doi:10.1007/s11627-021-10197-x

13. Rizzon C, Ponger L, Gaut BS. Striking similarities in the genomic distribution of tandemly arrayed genes in *Arabidopsis* and rice. PLOS Computational Biology 2006;2(9):e115, doi:10.1371/journal.pcbi.0020115

14. Yu J, Ke T, Tehrim S, et al. PTGBase: an integrated database to study tandem duplicated genes in plants. Database (Oxford) 2015;2015(doi:10.1093/database/bav017

15. Pan D, Zhang L. Tandemly arrayed genes in vertebrate genomes. Comp Funct Genomics 2008;2008(545269, doi:10.1155/2008/545269

16. Tuominen LK, Johnson VE, Tsai CJ. Differential phylogenetic expansions in BAHD acyltransferases across five angiosperm taxa and evidence of divergent expression among *Populus* paralogs. BMC Genomics 2011;12(236

17. Hanada K, Zou C, Lehti-Shiu MD, et al. Importance of lineage-specific expansion of plant tandem duplicates in the adaptive response to environmental stimuli. Plant Physiol 2008;148(2):993–1003, doi:10.1104/pp.108.122457

18. Baumgarten A, Cannon S, Spangler R, et al. Genome-level evolution of resistance genes in *Arabidopsis thaliana*. Genetics 2003;165(1):309–319, doi:10.1093/genetics/165.1.309

19. Jander G, Barth C. Tandem gene arrays: a challenge for functional genomics. Trends in Plant Science 2007;12(5):203–210, doi:10.1016/j.tplants.2007.03.008

20. Qi Y, Li X, Zhang Y, et al. Targeted deletion and inversion of tandemly arrayed genes in *Arabidopsis thaliana* using zinc finger nucleases. G3 Genes|Genomes|Genetics 2013;3(10):1707–1715, doi:10.1534/g3.113.006270

21. Lee HJ, Kim E, Kim JS. Targeted chromosomal deletions in human cells using zinc finger nucleases. Genome Res 2010;20(1):81–9, doi:10.1101/gr.099747.109

22. Richter J, Watson JM, Stasnik P, et al. Multiplex mutagenesis of four clustered CrRLK1L with CRISPR/Cas9 exposes their growth regulatory roles in response to metal ions. Sci Rep 2018;8(1):12182, doi:10.1038/s41598-018-30711-3

23. Seto D, Laflamme B, Guttman DS, et al. The *Arabidopsis* ZED1-related kinase genomic cluster is specifically required for effector-triggered immunity. Plant Physiol 2020;184(4):1635–1639, doi:10.1104/pp.20.00447

24. Čermák T, Curtin SJ, Gil-Humanes J, et al. A multipurpose toolkit to enable advanced genome engineering in plants. Plant Cell 2017;29(6):1196–1217, doi:10.1105/tpc.16.00922

25. Bewg WP, Harding SA, Engle NL, et al. Multiplex knockout of trichome-regulating MYB duplicates in hybrid poplar using a single gRNA. Plant Physiology 2022;189(516–526, doi:10.1101/2021.09.09.459666

26. Xue L-J, Guo W, Yuan Y, et al. Constitutively elevated salicylic acid levels alter photosynthesis and oxidative state, but not growth in transgenic *Populus*. Plant Cell 2013;25(2714–2730

27. Frost CJ, Nyamdari B, Tsai C-J, et al. The tonoplast-localized sucrose transporter in *Populus* (PtaSUT4) regulates whole-plant water relations, responses to water stress, and photosynthesis. PLoS One 2012;7(8):e44467

28. Vaser R, Sović I, Nagarajan N, et al. Fast and accurate de novo genome assembly from long uncorrected reads. Genome Res 2017;27(5):737–746, doi:10.1101/gr.214270.116

29. Koren S, Walenz BP, Berlin K, et al. Canu: scalable and accurate long-read assembly via adaptive k-mer weighting and repeat separation. Genome Research 2017;27(5):722–736

30. Thorvaldsdóttir H, Robinson JT, Mesirov JP. Integrative Genomics Viewer (IGV): high-performance genomics data visualization and exploration. Briefings in Bioinformatics 2013;14(2):178–192, doi:10.1093/bib/bbs017

31. Dellaporta SL, Wood J, Hicks JB. A plant DNA minipreparation: Version II. Plant Mol Biol Rep 1983;1(4):19–21, doi:10.1007/BF02712670

32. Tsai C-J, Xu P, Xue L-J, et al. Compensatory guaiacyl lignin biosynthesis at the expense of syringyl lignin in *4CL1*-knockout poplar. Plant Physiology 2020;183(123–136, doi:10.1104/pp.19.01550

33. Xue L-J, Tsai C-J. AGEseq: Analysis of genome editing by sequencing. Molecular Plant 2015;8(9):1428–1430, doi:10.1016/j.molp.2015.06.001

34. Inglis PW, Pappas MCR, Resende LV, et al. Fast and inexpensive protocols for consistent extraction of high quality DNA and RNA from challenging plant and fungal samples for high-throughput SNP genotyping and sequencing applications. PloS one 2018;13(10):e0206085, doi:10.1371/journal.pone.0206085

35. Xue L-J, Alabady MS, Mohebbi M, et al. Exploiting genome variation to improve next-generation sequencing data analysis and genome editing efficiency in *Populus tremula* x *alba* 717-1B4. Tree Genetics & Genomes 2015;11(82, doi:10.1007/s11295-015-0907-5

36. Zhou X, Jacobs TB, Xue L-J, et al. Exploiting SNPs for biallelic CRISPR mutations in the outcrossing woody perennial *Populus* reveals 4-coumarate:CoA ligase specificity and redundancy. New Phytologist 2015;208(298–301, doi:10.1111/nph.13470

37. Elorriaga E, Klocko AL, Ma C, et al. Variation in mutation spectra among CRISPR/Cas9 mutagenized poplars. Frontiers in Plant Science 2018;9(594), doi:10.3389/fpls.2018.00594

38. Bortesi L, Zhu C, Zischewski J, et al. Patterns of CRISPR/Cas9 activity in plants, animals and microbes. Plant Biotechnol J 2016;14(12):2203–2216, doi:10.1111/pbi.12634

39. Wu Q, Shou J. Toward precise CRISPR DNA fragment editing and predictable 3D genome engineering. Journal of Molecular Cell Biology 2020;12(11):828–856, doi:10.1093/jmcb/mjaa060

40. Fleiss A, O’Donnell S, Fournier T, et al. Reshuffling yeast chromosomes with CRISPR/Cas9. PLOS Genetics 2019;15(8):e1008332, doi:10.1371/journal.pgen.1008332

41. Muramoto N, Oda A, Tanaka H, et al. Phenotypic diversification by enhanced genome restructuring after induction of multiple DNA double-strand breaks. Nature Communications 2018;9(1):1995, doi:10.1038/s41467-018-04256-y

42. Xiao A, Wang Z, Hu Y, et al. Chromosomal deletions and inversions mediated by TALENs and CRISPR/Cas in zebrafish. Nucleic Acids Research 2013;41(14):e141–e141, doi:10.1093/nar/gkt464

43. Choi PS, Meyerson M. Targeted genomic rearrangements using CRISPR/Cas technology. Nature Communications 2014;5(1):3728, doi:10.1038/ncomms4728

44. Zhou H, Liu B, Weeks DP, et al. Large chromosomal deletions and heritable small genetic changes induced by CRISPR/Cas9 in rice. Nucleic Acids Research 2014;42(17):10903–10914, doi:10.1093/nar/gku806

45. Schmidt C, Pacher M, Puchta H. Efficient induction of heritable inversions in plant genomes using the CRISPR/Cas system. The Plant Journal 2019;98(4):577–589, doi:https://doi.org/10.1111/tpj.14322

46. Beying N, Schmidt C, Pacher M, et al. CRISPR–Cas9-mediated induction of heritable chromosomal translocations in *Arabidopsis*. Nature Plants 2020;6(6):638–645, doi:10.1038/s41477-020-0663-x

47. Simsek M, Adnan H. Effect of single mismatches at 3’-end of primers on polymerase chain reaction. J Sci Res Med Sci 2000;2(1):11–4

48. Lee H, Schatz MC. Genomic dark matter: the reliability of short read mapping illustrated by the genome mappability score. Bioinformatics 2012;28(16):2097–105, doi:10.1093/bioinformatics/bts330

49. Schmitz RJ, Grotewold E, Stam M. Cis-regulatory sequences in plants: Their importance, discovery, and future challenges. Plant Cell 2022;34(2):718–741, doi:10.1093/plcell/koab281

