## Supplemental Data for "Multiplex editing of the *Nucleoredoxin1* tandem array in poplar: from small indels to translocations and complex inversions" (from another version of this preprint: version 1, 2022-08-27)

**Table S1.** Primers used in this study.

| Target | Pr | Primer sequence (5' to 3') | Purpose and note |
|---|---|---|---|
| U6-gRNA | F | GTTTTCCCAGTCACGACGTTGTA | CRISPR vector construction |
| | R | TGGCTTGCTATTCCCTTCTCAAGCCTACTGGTTCGCTTGA | |
| gRNA-SF | F | GAGAAGGGAATAGCAAGCCAGTTTTAGAGCTAGAAATAGCAAGTT | CRISPR vector construction |
| | R | CGCACAGATGCGTAAGGAGAAA | |
| All PtaNRX1s | F | GCAATTGTTCTAGCMTTTTGAGT | Multigene PCR, Figs. 1C, 2E, 3B |
| | R | AAGTTCGTGYTTCACTTTCTCTG | |
| All PtaNRX1s | F | TTGGCAATTGTTCTAGCCTTT | Multigene PCR, Fig. 2C |
| | R | GATGYTCCTCGGTAAATGGAA | |
| All PtaNRX1s | F | CCTACACGACGCTCTTCCGATCTGAGGTGGTSTTCRTTTCTTCTGA | Amplicon sequencing Set 1 |
| | R | GTTCAGACGTGTGCTCTTCCGATCCCTCTTACTTTGAAYAMTTYCT | |
| All but PtaNRX1.2 | F | CCTACACGACGCTCTTCCGATCTGARKTGGTBTTCATTTCTTCTGA | Amplicon sequencing Set 2 |
| | R | GTTCAGACGTGTGCTCTTCCGATCCATCCACRCCATGYTC | |
| PtaNRX1.2 | F | CCTACACGACGCTCTTCCGATCTGARKTGGTBTTCATTTCTTCTGA | Amplicon sequencing Set 3 |
| | R | GTTCAGACGTGTGCTCTTCCGATCTGGCCAATTATGACAAGATTAGGG | |
| All PtaNRX1s | F | CCTACACGACGCTCTTCCGATCTGGGABCCCTCTTACTTYGAA | Amplicon sequencing Set 4 |
| | R | GTTCAGACGTGTGCTCTTCCGATCGCAATTTGGTTGGGAAGATTGT | |
| PtaNRX1.1 | F | GARKTGGTBTTCATTTCTTCTGA | Gene-specific PCR |
| | R | CATGCGACGTTAGCAGAG | |
| PtaNRX1.2 | F | TGATTCGTTAAGGGCTGATGTT | Gene-specific PCR, also amplify 1.1/1.3/1.4/1.5 with 1 to 2 mismatches |
| | R | TGGCCAATTATGACAAGATTAGGG | |
| PtaNRX1.3 | F | TGATTCGTTAAGGGCTGATGTT | Gene-specific PCR, also amplify 57, 53, 47, and 17 fusions |
| | R | TCACAGCACATCGTATGGATT | |
| PtaNRX1.4 | F | CTTGTCTTATTATAAGTTGACGTCTT | Gene-specific PCR |
| | R | ATGCAAGGTTAGCATAAAACAAG | |
| PtaNRX1.5 | F | TTTATGATTAAGAAAAAGCTCCCT | Gene-specific PCR |
| | R | ATCTGCCATTACAGATAGCAAG | |
| PtaNRX1.6 | F | TTTGTTTACCCTCTTRATTTAWGGG | Gene-specific PCR |
| | R | CACATCCKATGASCATRSGCTG | |
| PtaNRX1.7 | F | TTTGTTTACCCTCTTRATTTAWGGG | Gene-specific PCR |
| | R | TCACAGCACATCGTATGGATT | |
| PtaNRX1p$_{c10.1}$ | F | CGGAGACTCGCAAACTTCTT | Gene-specific PCR |
| | R | GGGATCTGCCACTTCTAATAGC | |
| PtaNRX1.7-1.2 & 1.6-1.2 fusions | F | TTTGTTTACCCTCTTRATTTAWGGG | Fusion-specific, also amplify PtaNRX1.6 and PtaNRX1.7 with 2 mismatches |
| | R | TGGCCAATTATGACAAGATTAGGG | |

**Table S2.** xGen lockdown probes used in NRX1-capture sequencing

| Probe Name | Sequence |
|---|---|
| probe_01 | ATAGCATTTGGAATCGGCAATTGTTCTAGCCTTTTGAGTAGTTCATCACTTAACAATGGCCAACGAAGACGTTTCGCACGACCTTTCATCGCTTCTTTCGTCGGAAGAGAGGGACTTTCT |
| probe_02 | AAGAGAGGGACTTTCTCATCCGCAACAATGGTGACCAGGTACTGCTTTTAGCTTATGGATTTCTGTTTTTCTCGTAATTAAGATGGATATCATGTTTAAATTAGACTGCGAACCTTGCTG |
| probe_03 | ACTGCGAACCTTGCTGAAGAACAAGTTTATGACCAATTTCTAGAGCTTTTATGATTAAGAAAATCCTCAATTTTTGTTTACCCTCTTAATTTAAGGGCTAATGTTTTTTCTGATTATGCT |
| probe_04 | TTTTTCTGATTATGCTCCACGCTAATTTTAGGTTAGGGTCAGCAATTTGGTTGGGAAGATTGTGGGATTCTATTTCTCTGGTTATTGGTGCGGGCCGTGCCGTAATTTCACTCCGTTGTT |
| probe_05 | ATTTCACTCCGTTGTTGGTAGAAGTCTATGAACAGCTATCATCCAAAGGGGACTTTGAGGTGGTCTTCATTTCTTCTGACGGACACGATGAATTCTTCAACACATACTTCTCCGAAATGC |
| probe_06 | TGATACGGAGACCCGCCAACGTCTTAAGAAAGTGTTCGAAGTAAGAGGGATCCCTAATCTTGTCATTTTTGATACGAATGGCAAGGTTTCATGCGATGATGGAGTCAGCACTGTCATGGA |
| probe_07 | CATGGAACATGGCGTGGATGGGTATCCGTTCAACCTTGATAGACTGATTTTCCTGAAAGAGCAAGAAGAGAAAGCTAAGAAGAATCAAACCATAAGCTCTATCTTGGTTTCCAGCTCACG |
| probe_08 | CTCACGTGATTATGTGATTTCAAATGATGGAAAAAAGGTAGATGGTTATTCAATTAGCTAGTTTTGAAATTGAATTGATGATGCCATAATTCACATTTTTTCTATCTGTAATGGCAGATC |
| probe_09 | CAGATCCCTTTGTTGGACCTTGAAGGGAAATTGGTTGGCTTGTATTTTTCAATCCATACGATGTGCTGTGAATTCACTCCAAAATTAGTGGAATTGTATAAGACGCTCAAGGAAAGAAGA |
| probe_10 | AGAAGAGAGAACTTTGAAGTAGTCCTAATATCTCTAGACGACGAAGAAGAAGACTTCAAAGAGAGTTTTGAGACAATGCCTTGGTTGGCATTGCCTTATAAGGACAAGAGCTGCGAGAAG |
| probe_11 | GAGAAGCTAGTGCGGTATTTTGAACTTAGAACCATTCCTAATCTTGTCATGATTGGCCAAGATGGGAAGACTTTGAACCCAAATGTAGCTGAACTGATCGAAGAACATGGTATTGAAGCC |
| probe_12 | GAAGCCTACCCATTTACACCGGGAAAGCTTGACGAGCTAGCTGCAATTGAAAAGGCAAAACTGGAATCGCAGACGCTTGAGTCAGTTTTGGTTAATGGGGAAAATGATTTTGTGATTGAC |
| probe_13 | ATTGACAAAAGTGGCTCCAAGGTAATTGTTTTTGTTCTATGCTAACCTCGCATGTTGTTACCGCATTACTTTCAATAACTTTCTTATGATCACGAATTGAATTCATTAGCAAAGGGTTTT |
| probe_14 | GGTTTTTTTTTTTTTTATCATTCAAATGGTGTGCTCTTGATTAGGTCCCAGTGTCTGAACTAGTTGGAAAGAACATTCTTCTTTACTTCTCAGCTCAATGGTGCCCTCCATGTCGTGCCT |
| probe_15 | GTGCCTTTTTACCCAAGCTAATTGAAGCATACCACACAATTAAAAGAAAAGACAATGCATTTGAGGTGATCTTCATCTCAAGTGACAGAGATCAATCCACCTTTGACGAGTTCTATTCAG |
| probe_16 | ATTCAGAAATGCCTTGGTTAGCCCTTCCATTTGGTGATGAAAGGAAACAAATCCTGAGTCGGAAATTCAAAATTCAAGGCATTCCTGCAGCTGTAGCGATTGGGCCAAGTGGCCGGACCA |
| probe_17 | GGACCATTACCAAGGAAGCTCGGATGCACCTGACAGCTTACGGGGCAGATGCTTTTCCATTTACCGAGGAACATCTAAAGCAATTGGAGGAGGAGCTTGAGGAAAAGGCAAAGGGGTGGC |
| probe_18 | GGTGGCCAGAGAAAGTGAAACACGAACTTCATACTGAGCATGAGCTGTTACGTAGTAAACGCAAAACATATATTTGCGATGGCTGTGGGGAAACAGGAAATAGTTGGTCTTTCCATTGCA |
| probe_19 | ATTGCAAAGAATGTGACTTTGATCTTCACCCCAAGTGTGCTTTGAAGGAAGATGAAGATACTGGAAGTGAAAAGGGAAATGAAGGATGGAACTGCGATGGAGATGTGTGCCGCCGAGCCT |
| probe_20 | GAGCCTAAGCAGGCAAGCGCTCCTTGTATCTGGTTTTTTGGGCTTTCTGCTGTAATAAGCGTGCGATGTAAGGTGTTTGTTGCATGTGTGTGAAGATGTGGATGTCTTTCCTGTTATTGA |
| probe_21 | TATTGATATGTAATTAAGAGGTGTGTTGCTTTAGGATTTTCATGTCCTGTATTTCGATACGAAGAATAAATAACCGGATTGGCTAGGTTTATGTGAATCCTGTGACCCTTTTGAAAC |
| probe_22 | CTGGAAGAGGTGCTATGCATGGTTCCATTGTGCCACTCTATTTGTTTCTTTTTGGCTGGATGAAGCAGCAGCTTCACCTCTGGTGCAAATATTCCTGTAATTGTTCTGCAGATTAAAAAA |
| probe_23 | AAAAGATGGCTGATTGTGACTAAACCTACATTATATAATGAAACCTTTCTAACTCCACCGTAAAAGAAACAAGAATCTCCGATCCAAGCAACTGGATAGTCTCTTTTTTCTCTCTCTCTG |
| probe_24 | TGCCTTCTCACATTACCTGTTGAGGCTGGTTTCAGAATCAAGAAAGCGGCATTAATCTTAACTCAGAACTAAGCAAGAAAAACTCATCCAATAGAAATGGCAATTCTTCTGTTGCTGATG |
| probe_25 | TGACTGGTTGACAAATGAAACAACAGCAAGTGGAAACTGGGTTCATTACAGTTATTTGAATGGCGCTAAGGGTGGAACCAAAATCACGAGCAGGCTATTGCTTGTACCAGTCGATGGGAG |
| probe_26 | AGGCAAAGGACGGGTAGGATTAATGGCGAGGTTCTAGAAATTGGTGAAGCCAACAGCTGTACCAAGAACACTGAGATGATGGTAAGAGCTTATTTGTTATTCTGTTAATGTTTTTTTTTC |
| probe_27 | TCTTTTAAGTGTTTTTAAATTGAAAATATTTTAAAATAATATTTTTTATTTTTTAAAATTTATTTCTCATTTCAACGTATCAAAATAAAAGTAATCTAAAACAAAATAAAAATTTATTTT |
| probe_28 | TTAAAATATTATTAAAAAATAATTTTAAAACACTAAAAATAATGTAATCTAAAACAATTTTTTTCGAAGATTTTTTTCAATCTAAAAAACAACATGCCTAAATAATTTACTTTACTTGTG |
| probe_29 | TGTGTGTTTAGTTTTAAAATTCAATTCAGCTAAATATGTTAACCTGAAGCTTGAACCAGGCCGTGTTTATAAAAAAATTAAAAAGGTTAAAAATGTCTCGTGTACTACAAAAGTTTTATC |
| probe_30 | TCTTTAAACTTTGTGTAATGATACGACTAGTAAAACCTGTCAAAACTTAATTATAATTATTTAACTTTATATTTTTCATAAAAAAAAAAAGTATTACAAGTTTAAAAAATATACTTGTTA |
| probe_31 | TACCAAAAAAAAAACTCAAATGAGTTAACTTTGTTTGCAGTGAGAGTAAGAATTTTTGTTATATATATATATATATATTCAAAGAAACTACAAAAAGGTAAGGTTATGAAATAAAAAGGC |
| probe_32 | GCAAGTTAGAAAGTGCAGAAGAAATTTTCTTGTTAGTTTTTAAATTTTATATTGTAGAATTTATGTCTACCAATTCATATTTAATAATAAAAATTAACCCGTCCCTAGATTTACTTGATA |
| probe_33 | TATTTAATTTTAAAATTTAAAATAAATAATTAAGGTGCACGCTACATCGGTTACTTTAAAAAACATTTGAAAGGGTGTGTATCCTAATCTCCACCGTTAATCATACCGTCCATGAGTCTG |
| probe_34 | TGATCGCACGGTTTTTATAACATCTTCAGTCTTCTGTCAGTATATTCCAATTTTGCAGATCAAATCATATCTCTCGCTTTCTCGGTAAGATTCTTCCTCTCTCTCTCCTTTACCATGCGA |
| probe_35 | GATCTGCGTTTAGAATTATTACTTCAATCCTTTCGTTAATTCCTTAATTATGATCTAGGGTTTTGGGGTGTTAATTAAATTGAATTATTATGTTTGGGTGATCGTTGTAATTTTTGGTGG |
| probe_36 | GGTTAGATGGATTGGCAAGGACAGAAACAAGCGGAGCTATGGATGCAGATACTGCTGTTAGTTTTCGCGGCGGCGGCTTTAGTTATAGGTTATATAATAGGATCGTTTCGAATGATGATG |
| probe_37 | TGCTAATTTACGCCGGTGGAGTGGTTTTCACCTCACTGGTTACTGTCCCTAATTGGCCCTTCTTCAATCGCCATCCTCTCAAGTGGTTGGACCCAAGCGAAGCTGAGAAGCACCCTAAGC |
| probe_38 | GCCACAGAAGGCTGTGGTTTTCAAGGACAAGAAGAAATCGTCCAAGAAGTAGGTTGGTTTTGTTAAATTTGACTGGTTTTTTAGTCTAAGCATTTGTTTGCTAGGGTTTTTAAATCTTCA |
| probe_39 | CAATTTAATGGATGGAATGAAACGTTATGGGATGAATTGGGTGTTAGAATAACTTTGTTTTGCTATAAATTTTATACCTTTTAGACAATGCCATTCGAGTTTCAGTTAGTTTTTTTAACT |
| probe_40 | GCATGTTTGAAGTGTTGGTTCTCGTTTATAGCTGTATTGATTTCTATTCATTGATGCTCGGACCATTGTTTCTGATTAGCTTTGCCCTCTCAATTGCTTTAAGTACTTCATTTTAAAAAA |
| probe_41 | TTAAAAAACCAGTTATGGATGATCTTGATTTTAATTATGTTTCCTCCATTGATAGATTTTCTAAGCTTGTTTTGACATGTGCATCCTTTTTAATGTCATGCAGAATACAAGAGAGACAGC |
| probe_42 | GAGACAGCGTTTGCTCTAAGGAAGTTGCCTCTGGTCAAGGCCAAGAGGTATTTGGAAGATGTTATGGCTCACAAACAGGCCATTCCATTCCGGCGTTTCTGTGGTGGAGTTGGGCGTACT |
| probe_43 | GGCGTACTGCACAAGCAAAGAACAGACACTCTAACGGACAAGGAAGATGGCCTGCCAAGTCTGCCAAGTTCATCCTGGATTTACTCAAGAATGCCGAGAGCAATGCTGAGGTATATCTAT |
| probe_44 | ATATCTATCCTCTTATTTGTCTTCACTTTCTCATCAACTTCTGCTTGCCATTTTACTGACATTACTTACCTTTCCAGGTCAAGGGTTTGGATGTGGATGCACTCTACATTTCTCACATCC |
| probe_45 | TCACATCCAGGTGAATCAGGCACAGAAGCAGAGACGTCGTACTTACCGGGCGCATGGAAGAATTAATCGTAAGCTTGTTTTGATTAGATTGGTATTAGTTAATCTTACAATTACATTTCC |
| probe_46 | ACATTTCCATTTGTATTCACAGGACTCTTTTATTTTCCTTTCCAGCTTACATGTCCAGCCCTTGCCACATTGAGTTGACTTTATCTGAAAAGGAAGAGCCAGTTAAGAAAGAGGTAATTC |
| probe_47 | GGTAATTCATTTCTTTCTAATTGGGTAGATGGTTTAATTAATATACCAGTGTTTTTTGGACCTTTCTAATTCTGTGTACTTGTATTGCAGCCTGAGACCCAGCTAGCAACCAGCAAGTCA |
| probe_48 | GCAAGTCAAAGAAGTCTCAGGTTTCCTCTTGAGTGGCGATCTTAGTTCAGTGTGGATCCCTTTTTGATTTCTGTGTTTTTTTCTTAAGAAGGATTTTGTTGCGTATCTTATAGAATGATC |
| probe_49 | GAATGATCAGATTTTTAGATGTACTGGACATCTTGTATGTTGATTTCGAATTATGTTTAGTTCTTTGTTTTATGAGAAATGTCTTGGCGAGTTTATTTGAAGTCTATTTTCATCACAGTT |

**A.** *P. trichocarpa*

| | *PotriNRX1.1* | *PotriNRX1.2* | *PotriNRX1.3* | *PotriNRX1.4* | *PotriNRX1.5** | *PotriNRX1.6* | *PotriNRX1.7* | *PotriNRX1.8* |
|---|---|---|---|---|---|---|---|---|
| *PotriNRX1.1* | | | | | | | | |
| *PotriNRX1.2* | 99% | | | | | | | |
| *PotriNRX1.3* | 96% | 96% | | | | | | |
| *PotriNRX1.4* | 95% | 95% | 96% | | | | | |
| *PotriNRX1.5** | 89% | 89% | 93% | 90% | | | | |
| *PotriNRX1.6* | 95% | 95% | 96% | 98% | 90% | | | |
| *PotriNRX1.7* | 95% | 95% | 95% | 95% | 88% | 95% | | |
| *PotriNRX1.8* | 96% | 95% | 97% | 99% | 90% | 98% | 96% | |

**B.** *P. tremula x alba*

| | *PtaNRX1.1* | *PtaNRX1.2** | *PtaNRX1.3* | *PtaNRX1.4* | *PtaNRX1.5* | *PtaNRX1.6* | *PtaNRX1.7* |
|---|---|---|---|---|---|---|---|
| *PtaNRX1.1* | | | | | | | |
| *PtaNRX1.2** | 56% | | | | | | |
| *PtaNRX1.3* | 95% | 69% | | | | | |
| *PtaNRX1.4* | 99% | 69% | 96% | | | | |
| *PtaNRX1.5* | 98% | 49% | 94% | 98% | | | |
| *PtaNRX1.6* | 94% | 68% | 95% | 94% | 93% | | |
| *PtaNRX1.7* | 93% | 69% | 96% | 94% | 92% | 96% | |

**Figure S1.** Coding sequence identity matrix among poplar *NRX1* TAGs. A, *PotriNRX1s* from *P. trichocarpa.* B, *PtaNRX1s* from *P. tremula* x *alba* INRA 717-1B4. Asterisks indicate sequence truncation or alignment gaps.

```
PtaNRX1.3      GCAATTGTTCTAGCCTTTTGAGTAG-TTCATCACATAGCTAGCTATGGCCACCGAAGACG
PtaNRX1.4      GCAATTGTTCTAGCCTTTTGAGTAG-TTCATCACAGAGCTAGCTATGGCCACCGAAGACG
PtaNRX1.1      GCAATTGTTCTAGCCTTTTGAGTAG-TTCATCACAGAGCTAGCTATGGCCACCGAAGACG
PtaNRX1.5      GCAATTGTTCTAGCCTTTTGAGTAGTTTCATCACATAGCTAGCTATGGCCACCGAAGACG
PtaNRX1.2      GCAATTGTTCTAGCCTTTTGAGTAG-TTCATCACATAGCTAGCTATGGCCACCGAAGACG
PtaNRX1.7      GCAATTGTTCTAGCCTTTTGAGTAG-TTCATCAC----TTAACAATGGCCAACGAAGACG
PtaNRX1.6      GCAATTGTTCTAGCCTTTTGAGTAG-TTCATCAC----TTAACAATGGCCAACGAAGACG
               ************************* ********     ** * ******* ********

PtaNRX1.3      TTTCTCTCGACCTTTCAACGCTTCTTTCTTCAGAAGAGAGGGACTTTCTCATCCGCAACA
PtaNRX1.4      TTTCTCTGGACCTTTCAACGCTTCTTTCTTCAGTAGAGAGGGACGTTCTCATCCGCAACA
PtaNRX1.1      TTTCTCTCGACCTTTCAACGCTTCTTTCTTCAGAAGAGAGGGACGTTCTCATCCGCAACA
PtaNRX1.5      TTTCTCTCGACCTTTCGACGCTTCTTTCTTCAGCAGAGAGGGACTTTCTCATCCGCAACA
PtaNRX1.2      TTTCTCTCGACCTTTCAACGCTTCTTTCTTCAGCAGAGAGGGACTTTCTCATCCGCAACA
PtaNRX1.7      TTTCGCACGACCTTTCATCGCTTCTTTCGTCGGAAGAGAGGGACTTTCTCATCCGCAACA
PtaNRX1.6      TTTCGCACGATCTTTCATCCCTTCTTTCGTCGGAAGAGAGGGACTTTCTCATCCGCAACA
               **** *  ** *****  * ******** ** * ********** ***************

PtaNRX1.3      ATGGTGATCAGGTACTGCTTCTCTGATCTTATTCAAGTCTTCTTCTTGTCTTATTATAAG
PtaNRX1.4      ATGGTGATCAGGTACTGCTTCTCTGATCTTATTCAAGTCTTCTTCTTGTCTTATTATAAG
PtaNRX1.1      ATGGTGATCAGGTACTGCTTCTCTGATCTTATTCAAGTCTTCTTCTTGTCTTATTATAAG
PtaNRX1.5      ATGGTGATCAGGTACTGCTTCTCTGATCTTATTCAAGTCTTCTTCTTGTCTTATTATAAG
PtaNRX1.2      ATGGTGATCAGGTACTGCTTCTCTGATCTTATTCAAGTCTTCTTCCTGTCTTATTATAAG
PtaNRX1.7      ATGGTGACCAGGTACTGCTTTT---AGCTTAT----GGATTTCT----------------
PtaNRX1.6      ATGGTGACCAGGTACTGCTTTT---AGCTTAT----GGATTTCTCTGGTCTTAT-----G
               ******* ************ *   * *****    *  **  *

PtaNRX1.3      TTGA-------TAATTTTTATA-TGAACTTGTAGGTTTTTCTCGTAATTAAGATGAATAT
PtaNRX1.4      TTGACGTCTTTTAATTTTTATA-TGAACTTGTAGGTTTTTCTCGTAATTAGGATGAATAT
PtaNRX1.1      TTGACGTCTTTTAATTTTTATA-TGAACTTGTAGGTTTTTCTCGTAATTAGGATGAATAT
PtaNRX1.5      TTGACGTCTTTTAATTATTATA-TGAACTTGTAGGTTTTTCTCGTAATTAAGATGAGTAT
PtaNRX1.2      TTGACGTCTTTTAATTTTTATA-TGAACTTGTAGGTTTTTCTCGTAATTAAGATGAATAT
PtaNRX1.7      ----------------------------------GTTTTTCTCGTAATTAAGATGGATAT
PtaNRX1.6      CGATCTACTTCTACTTGTCTTATTAAACTTGCAGGTTTTTCTCGTAATTAGGATGGATAT
                                                 **************** ****  ***

PtaNRX1.3      CATGTTTAAATTAGACTGCAAGCCTTGCTGAAGAACAAATTTATGACCAATTTCTGAAGC
PtaNRX1.4      CATGTTTAAATTAGACTGAAAGCCTTGCTGAAGAACAAATTTATGACCAATTTCTGAAGC
PtaNRX1.1      CATGTTTAAATTAGACTGAAAGCCTTGCTGAAGAACAAATTTATGACCAATTTCTGAAGC
PtaNRX1.5      CATGTTTAAATTAGACTGTTAGCCTTGCTTAAGAACAAATTTATGACCAATTTCTGAAGC
PtaNRX1.2      CATGTTTAAATTAGACTGGTAGCCTTGCTTAAGAACAAATTTATGACCAATTTCTGAAGC
PtaNRX1.7      CATGTTTAAATTAGACTGCGAACCTTGCTGAAGAACAAGTTTATGACCAATTTCTAGAGC
PtaNRX1.6      CATGTTTAAATTAGACTGCGAACCTTGCTCAAGAACAAGTTTATGACCAATTTCTAGAGC
               ******************  * ******* ******** ****************  ***

PtaNRX1.3      TTTTTAATGATTAAGAAAAAGCTCCTTTTTTTGTGTTCCTTTTCTCACTGGCAAGAGAAA
PtaNRX1.4      TTTTTTATGATTAAGAAAAAGCTCCTTTTTTTTTGTTGCTTCTCTCACTGGCAAGAGAAA
PtaNRX1.1      TTTTTTATGATTAAGAAAAAGCTCCTTTTTTTTTGTTGCTTCTCTCACTGGCAAGAGAAA
PtaNRX1.5      -TTTTTATGATTAAGAAAAAGCTCCCTTTTTTTTGTTGCTTCTCTCACTGGCAAGAGAAA
PtaNRX1.2      -TTTTTATGATCAAGAAAAAGCTCCCTTTTTTTTGTTGCTTCTCTCACTGGCAAGAGAAA
PtaNRX1.7      --TTTTATGATTAAGAAAA---TCCTCAATTTTTGTTTACCCTCTTAAT-----------
PtaNRX1.6      --TTTTATGATTCAGAAAA---GCCTCAATTTTTGTTTACCCTCTTGAT-----------
                 *** *****  ******    **    *** ****     ***   *

PtaNRX1.3      GAAACCATGAATAAGATTTTGATTCGTTAAGGGCTG--ATG-TTTTTTTCTGATAATGTT
PtaNRX1.4      GAAACCATGAATAAAATTGTGATTCGTTAAGGGCTG--ATG-TTTTTTTCTGATAATGTT
PtaNRX1.1      GAAACCATGAATAAAATTGTGATTCGTTAAGGGCTG--ATG---TTTTTCTGATAATGTT
PtaNRX1.5      GAAACCATGAATAAAATTTTGATTCGTTAAGGGCTG--ATGTTTTTTTTCTGATAATATT
PtaNRX1.2      GAAACCATGAATAAAATTTTGATTCGTTAAGGGCTG--ATG-TTTTTTTCTGATAATGTT
PtaNRX1.7      --------------------------TTAAGGGCTA--ATG--TTTTTCTGATTATGCT
PtaNRX1.6      --------------------------TTATGGGCTATTTTTTTTTTTTTCTGATTATGCT
                                         *** *****    *    ********** **  *

PtaNRX1.3      CGAGATTAATTTTTAGGTTAAGGTCAGCAATTTGGTTGGGAAGATTGTGGGCTTCTATTT
PtaNRX1.4      CGAGATTAATTTTTAGGTTAAGGTCAGCAATTTGGTTGGGAAGATTGTGGGGTTCTACTT
PtaNRX1.1      CGACATTAATTTTTAGGTTAAGGTCAGCAATTTGGTTGGGAAGATTGTGGGGTTCTATTT
PtaNRX1.5      CGAGATTAATTTTTAGGTTAAGGTCAGCAATTTGGTTGGGAAGATTGTGGGGTTCTACTT
PtaNRX1.2      CGAGATTAATTTTTAGGTTAAGGTCAGCAATTTGGTTGGGAAGATTGTGGGGTTCTACTT
PtaNRX1.7      CCACGCTAATTTT-AGGTTAGGGTCAGCAATTTGGTTGGGAAGATTGTGGGATTCTATTT
PtaNRX1.6      CGACACTAATTTTAAGGTTACGGTCAGCAATTTGGTTGGGAAGATTGTGGGATTCTATTT
               * *   ******* ****** ****************************** ***** **
```

```
PtaNRX1.3      CTCTGGTTCGTGGTGCGGGCCGTGCCGTAATTTCACTCCATTGTTGGTAGAAGTCTATGA
PtaNRX1.4      CTCTGGTTCGTGGTGCGGCCCATGCCGTAATTTCACTCCTTTGTTGGTAGAAGTCTATGA
PtaNRX1.1      CTCTGGTTCGTGGTGCGGCCCATGCCGTAATTTCACTCCTTTGTTGGTAGAAGTCTATGA
PtaNRX1.5      CTCTGGTTCTTGGTGCGGCCCATGCCGTAATTTCACTCCTTTGTTTGTAGAAGTCTATGA
PtaNRX1.2      CTCTGGTTCGTGGTGCGGCCCATGCCGTAATTTCACTCCTTTGTTGGTAGAAGTCTATGA
PtaNRX1.7      CTCTGGTTATTGGTGCGGGCCGTGCCGTAATTTCACTCCGTTGTTGGTAGAAGTCTATGA
PtaNRX1.6      CTCTGGTTCGTGGTGCGGGCCGTGCCGCAATTTCACTCCGTTGTTGGTAGAAGTCTATGA
               ********  ******** ** ***** *********** ***** **************

PtaNRX1.3      ACAGCTATCATCCAAAGGGGACTTTGAGGTGGTCTTCATTTCTTCTGACGGAGACGATGA
PtaNRX1.4      ACAGCTATCATCCAAAGGCGACTTTGAGGTGGTGTTCATTTCTTCTGACAGAGATGATGA
PtaNRX1.1      ACAGCTATCATCCAAAGGCGACTTTGAGGTGGTGTTCATTTCTTCTGACAGAGATGATGA
PtaNRX1.5      ACAGCTATCATCCAAAGGGGACTTTGAGGTGGTCTTCATTTCTTCTGATAGAGATGATGA
PtaNRX1.2      ACAGCTATCATCCAAAGGGGACTTTGAGGTGGTCTTCATTTCTTCTGACAGAGATGATGA
PtaNRX1.7      ACAGCTATCATCCAAAGGGGACTTTGAGGTGGTCTTCATTTCTTCTGACGGACACGATGA
PtaNRX1.6      ACAGCTATCATCCAAAGGGGACTTTGAGGTGGTCTTCATTTCTTCTGACGGAGACGATGA
               ****************** ************** **************  ** * *****

PtaNRX1.3      ATCCTTCAACACATACTTCTCCGAAATGCCCTGGCTTGCTATTCCCTTCTCTGATACGGA
PtaNRX1.4      ATCCTTCAACACATATTTCTCCGAAATGCCCTGGCTTGCTATTCCCTTCTCTGATACGGA
PtaNRX1.1      ATCCTTCAACACATATTTCTCCGAAATGCCCTGGCTTGCTATTCCCTTCTCTGATACGGA
PtaNRX1.5      ATCCTTCAACACATACTTCTCCGAAATGCCCTGGCTTGCTATTCCCTTCTCTGATACGGA
PtaNRX1.2      ATCCTTCAACACATATTTCTCCGAAATGCCATGGCTTGCTATTCCCTTCTCTGATACGGA
PtaNRX1.7      ATTCTTCAACACATACTTCTCCGAAATGCCATGGCTTGCTATTCCCTTCTCTGATACGGA
PtaNRX1.6      ATCCTTCAACACATACTTCTCCGAAATGCCATGGCTTGCTATTCCCTTCTCTGATACGGA
               ** ************ ************** *****************************

PtaNRX1.3      GACCCGCAAACGTCTTAAGGAAGTGTTCAAAGTAAGAGGGATCCCTACTCTTGTCATTTT
PtaNRX1.4      GACCCGCAAACGTCTTAAGGAAGTGTTCAAAGTAAGAGGGGTCCCAAATCTTGTCATTTT
PtaNRX1.1      GACCCGCAAACGTCTTAAGGAAGTGTTCAAAGTAAGAGGGATCCCAAATCTTGTCATTTT
PtaNRX1.5      GACCCGCAAACGTCTTAAGGAAGTGTTCAAAGTAAGAGGGCTCCCTAATCTTGTCATTTT
PtaNRX1.2      GACCCGCAAACGTCTTAAGGAAGTATTCAAAGTAAGAGGGATC-----------------
PtaNRX1.7      GACCCGCCAACGTCTTAAGAAAGTGTTCGAAGTAAGAGGGATCCCTAATCTTGTCATTTT
PtaNRX1.6      GAGCCGCCAACGTCTTCAGGAATTGTTCAAAGTAAGAGGGATCCCTAAACTCGTCATTTT
               ** **** ******** ** ** * *** *********** **

PtaNRX1.3      TGATACGAATGGCAAGGTTTCATGCGATGATGGAGTCAGCACTGTCATGGAACATGGCGT
PtaNRX1.4      TGATGCGAATGGCAAGGTTTCCTGCGATGATGGAGTCAGCACTGTCAAGGAACATGGCGT
PtaNRX1.1      TGATGCGAATGGCAAGGTTTCCTGCGATGATGGAGTCAGCACTGTCAAGGAACATGGCGT
PtaNRX1.5      TGATGCGAATGGCAAGGTTTCCTGCGATGATGGAGTCAGCACTGTCAAGGAGCATGGCGT
PtaNRX1.2      ------------------------------------------------------------
PtaNRX1.7      TGATACGAATGGCAAGGTTTCATGCGATGATGGAGTCAGCACTGTCATGGAACATGGCGT
PtaNRX1.6      TGATGCTAATGGTAAGGTTTCCTGCGATAATGGAGTCAGACTTGTCAAGGAACATGGCGT

PtaNRX1.3      GGATGGGTATCCGTTCAACCTTGATAGACTGAATTTCCTGAAAGAGCAAGAAGAGAATGC
PtaNRX1.4      GGATGGGTATCCGTTCAACCTTGATAGACTGAATTTCCTGAAAGAGCAAGAAGAGAATGC
PtaNRX1.1      GGATGGGTATCCGTTCAACCTTGATAGACTGAATTTCCTGAAAGAGCAAGAAGAGAATGC
PtaNRX1.5      GGATGGGTATCCGTTCAACCTTGATAGACTGAATTTCCTGAAAGAGCAAGAAGAGAATGC
PtaNRX1.2      ------------------------------------------------------------
PtaNRX1.7      GGATGGGTATCCGTTCAACCTTGATAGACTGATTTTCCTGAAAGAGCAAGAAGAGAAAGC
PtaNRX1.6      GGATGGGTATCCGTTCAACCTTGATAGACTGAATTTCCTTAAAGAGCAAGAAGAGAATGC

PtaNRX1.3      TAAGAAGAATCAAACCATAAGCTCTATCTTGGTTTCCAGCTCACGTGATTATGTGATTTC
PtaNRX1.4      TAAGAAGAATCAAACCATAAGCTCTATCTTGGTTTCAAGCTCACGTGATTATGTGATTTC
PtaNRX1.1      TAAGAAGAATCAAACCATAAGCTCTATCTTGGTTTCAAGCTCACGTGATTATGTGATTTC
PtaNRX1.5      TAAGAAGAATCAAACCATAAGCTCTATCTTGGTTTCCAGCTCACGTGATTATGTGATTTC
PtaNRX1.2      ------------------------------------------------------------
PtaNRX1.7      TAAGAAGAATCAAACCATAAGCTCTATCTTGGTTTCCAGCTCACGTGATTATGTGATTTC
PtaNRX1.6      TAAGAAGAATCAAACCATAAGCTCTATCTTGGTTTCCAGCTCACGTGATTATGTGATTTC

PtaNRX1.3      AAATGATGGAAAAAAGGTAGATGTTTTCATTCAATTAGCTAGTTTTGAAATTGAAATGAT
PtaNRX1.4      AAATGATGGAAAAAAGGTAGATG-GTTCATTCAATTAGCTAGTTTTGAAATTGAATTGAT
PtaNRX1.1      AAATGATGGAAAAAAGGTAGATG-GTTCATTCAATTAGCTAGTTTTGAAATTGAATTGAT
PtaNRX1.5      TAATGATGGAAAAAAGGTAGATG-GTTCATTCAATTAGCTAGTTTTGAAATTGAATCGAT
PtaNRX1.2      ------------------------------------------------------------
PtaNRX1.7      AAATGATGGAAAAAAGGTAGATG-GTT-ATTCAATTAGCTAGTTTTGAAATTGAATTGAT
PtaNRX1.6      AAATGATGGAAAAAAGGTAGATG-TTTCATTCAATTAGCTAGTTTTGAAATTGAATTGAT
```

```
PtaNRX1.3       GATGCCATCATTCA--------CATTTTTGCTATCTGTAATGGCAGATCCCTGTGTTGGA
PtaNRX1.4       GATGCCATCATTCACTTTTTGCCTTTTTGGCTATCTGTAATGGCAGATCCCTGTGTTGGA
PtaNRX1.1       GATGCCATCATTCACTTTTTGCCTTTTTGGCTATCTGTAATGGCAGATCCCTGTGTTGGA
PtaNRX1.5       GATGCCATCATTCA--------CTTTCTTGCTATCTGTAATGGCAGATCCCTGTGTTGGA
PtaNRX1.2       ------------------------------------------------------------
PtaNRX1.7       GATGCCATAATTCA--------CATTTTTTCTATCTGTAATGGCAGATCCCTTTGTTGGA
PtaNRX1.6       GATGCCATAATTCA--------CATTTTTGCTATCTGTAATGTCAGATCCCTGTGTTGGA

PtaNRX1.3       GCTTGAAGGAAAATTGGTTGGCTTGTATTTTTCAATCCATAC------GATGTGCTGTGA
PtaNRX1.4       CCTTGAAGGAAAATTGGTTGGCTTGTATTTCTCAATCCATGCTCATAGGATGTGCCGTGA
PtaNRX1.1       CCTTGAAGGAAAATTGGTTGGCTTGTATTTCTCAATCCATGCTCACAGGATGTGCCGTGA
PtaNRX1.5       CCTTGAAGGAAAATTGGTTGGCTTGTATTTCTCAATCCATGCTCATAGGATGTGCCGTGA
PtaNRX1.2       ------------------------------------------------------------
PtaNRX1.7       CCTTGAAGGGAAATTGGTTGGCTTGTATTTTTCAATCCATAC------GATGTGCTGTGA
PtaNRX1.6       CCTTGAAGGAAAATTGGTTGGCTTGTATTTTTCAGCCCATGCTCATAGGATGTGCCGTGA

PtaNRX1.3       ATTCACTCCAAAACTAGTGGAATTGTATAAGACGCTCAAGGAAAAAAGAGAGAACTTTGA
PtaNRX1.4       ATTCACTCCTAAACTAGTTGAATTACACAAGCGGCTCAAGGAAAAAGGAGAGAACTTTGA
PtaNRX1.1       ATTCACTCCTAAACTAGTTGAATTACAAAAGCGGCTCAAGGAAAAAGGAGAGAACTTTGA
PtaNRX1.5       ATTCACTCCTAAACTAGTTGAATTATACAAGAGGCTCAAGGAAAAAGGAGAGAACTTTGA
PtaNRX1.2       ------------------------------------------------------------
PtaNRX1.7       ATTCACTCCAAAATTAGTGGAATTGTATAAGACGCTCAAGGAAAGAAGAGAGAACTTTGA
PtaNRX1.6       ATTCACTCCTAAACTAGTGGAATTGTATAAGACGCTCAAGGAAAGAAGAGAGAACTTTGA

PtaNRX1.3       AGTAGTCCTAATATCTCTAGACGACGAGGAAGAAGACTTCAAAGAGAGATTTGAGACAAT
PtaNRX1.4       AGTAGTCCTAATATCTCTAGACTTTGAGGAAAAACACTTCAAAGAGAGTTTTGAGACAAT
PtaNRX1.1       AGTAGTCCTAATATCTCTAGACTTTGAGGAAAAACACTTCAAAGAGAGTTTTGAGACAAT
PtaNRX1.5       AGTAGTCCTAATATCTCTAGACTTTGAGGAAAAACACTTCAAAGAGAGTTTTGAGACAAT
PtaNRX1.2       ------------------------------------------------------------
PtaNRX1.7       AGTAGTCCTAATATCTCTAGACGACGAAGAAGAAGACTTCAAAGAGAGTTTTGAGACAAT
PtaNRX1.6       AGTAGTCCTAATATCTCTAGACGACGAAGAAGAAGACTTCAAAGAGAGTTTTGAGACAAT

PtaNRX1.3       GCCTTGGTTGGCATTGCCATTTAAGGACAAGAGCTGCGAGAAGCTAGTGCGGTATTTTGA
PtaNRX1.4       GCCTTGGTTGGCATTGCCGTTTAAGGACAAGAGCTGCGAGAAGCTAGCGCGGTATTTTGA
PtaNRX1.1       GCCTTGGTTGGCATTGCCGTTTAAGGACAAGAGCTGCGAGAAGCTAGCGCGGTATTTTGA
PtaNRX1.5       GCCTTGGTTGGCATTGCCGTTTAAGGACAAGAGCTGCGAGAAGCTAGCGCGGTATTTTGA
PtaNRX1.2       ------------------------------------------------------------
PtaNRX1.7       GCCTTGGTTGGCATTGCCTTATAAGGACAAGAGCTGCGAGAAGCTAGTGCGGTATTTTGA
PtaNRX1.6       GCCTTGGTTGGCATTGCCGTTTAAGGACAAGAGCTGCGGGAAGCTAGTGCGGTATTTTGA

PtaNRX1.3       ACTTGGAACCATTCCTAATCTTGTCATAATTGGTCAAGATGGGAAGACTTTGAACCCAAA
PtaNRX1.4       ACTTAGAACCATTCCTAATCTTGTCATAATTGGCCAAGATGGGAAGACTTTGAACCCAAA
PtaNRX1.1       ACTTAGAACCATTCCTAATCTTGTCATAATTGGCCAAGATGGGAAGACTTTGAACCCAAA
PtaNRX1.5       ACTTAGAATCATTCCTAATCTTGTCATAATTGGCCAAGATGGGAAGACTTTGAACCCAAA
PtaNRX1.2       -------------CCTAATCTTGTCATAATTGGCCAAGATGGGAAGACTTTGAACCCAGA
PtaNRX1.7       ACTTAGAACCATTCCTAATCTTGTCATGATTGGCCAAGATGGGAAGACTTTGAACCCAAA
PtaNRX1.6       ACTTCGAACCATTCCTAAGCTTGTCATAATTGGCCAAGATGGGAAGACTTTGAACCCAAA
                             ***** ******** ***** ************************ *

PtaNRX1.3       TGTAGCTGAACTGATCGAAGAACATGGTATTGAAGCCTACCCATTTACACCGGAAAAGCT
PtaNRX1.4       TGTAGTTGAACTCATCGAAGACCATGGAATTGAAGCCTACCCATTTACACCAGAAAAGCT
PtaNRX1.1       TGTAGTTGAACTCATCGAAGACCATGGAATTGAAGCCTACCCATTTACACCGGAAAAGCT
PtaNRX1.5       TGTAGTTGAACTCATTGAAGACCATGGAATTGAAGCCTACCCATTTACACCGGAAAAGCT
PtaNRX1.2       TGTAGTTGAACTCATCGAAGACCATGGAATTGAAGCCTACCCATTTACACCGGAAAAGCT
PtaNRX1.7       TGTAGCTGAACTGATCGAAGAACATGGTATTGAAGCCTACCCATTTACACCGGAAAAGCT
PtaNRX1.6       TGTAGCTGAACTGATCGAAGAACATGGTATTGAAGCCTACCCATTTACACCGGAAAAGCT
                ***** ****** ** ***** ***** ********************** ********

PtaNRX1.3       TGACGAG-CTAGCTGCAATTGAAAAGGCAAAACTGGAATCGCAGACGCTTGAATCAGTTT
PtaNRX1.4       TGATGAG-CTAGCTGCAATTGAAAAGGCAAAACTGGAATCGCAGACGCTTGAATCAGTTT
PtaNRX1.1       TGAAGAG-CTAGCTGCAATTGAAAAGGCAAAACTGGAATCGCAGACGCTTGAGTCAGTTT
PtaNRX1.5       TGAAGAGCCTAGCTGCAATTGAAAAGGCAAAACTGGAATCGCAGACGCTTGAGTCAGTTT
PtaNRX1.2       TGATGAG-CTAGCTGCAATTGAAAAGGCAAAACTGGAATCGCAGACGCTTGAGTCAGTTT
PtaNRX1.7       TGACGAG-CTAGCTGCAATTGAAAAGGCAAAACTGGAATCGCAGACGCTTGAGTCAGTTT
PtaNRX1.6       TGACGAG-CTAGCTGCAATTGAAAAGGCAAAACTGGAATCGCAGACGCTTGAGTCAGTTT
                *** *** ******************************************** *******
```

```
PtaNRX1.3       TGGTAAATGGGGAAAATGATTTTGTGATTGACAAAAGTGGCTCCAAGGTAATT-TTCTTG
PtaNRX1.4       TGGTTAATGGGGAAAATGATTTTGTGATTGACAAAAGTGGCTCCAAGGTAATT-TTCTTG
PtaNRX1.1       TGGTTAATGGGGAAAATGATTTTGTGATTGACAAAAGTGGATCCAAGGTAATT-TTTTTG
PtaNRX1.5       TGGTTAATGGGGAAAATGATTTTGTGATTGACAAAAGTGGATCCAAGGAAATT-TTTTTG
PtaNRX1.2       TGGTTAATGGGGAAAATGATTTTGTGATTGACAAAAGTGGATCCAAGGTAATT-TTTTTG
PtaNRX1.7       TGGTTAATGGGGAAAATGATTTTGTGATTGACAAAAGTGGCTCCAAGGTAATTGTTTTTG
PtaNRX1.6       TGGTTAATGGGGAAAATGATTTTGTGATTGACAAAAGTGGCTCCAAGGTAATTGTTTTTG
                **** ********************************** ******* **** ** ***

PtaNRX1.3       TTTTATGCTAACCTCGCATGTTGTTATTGCATTACTTTCAATAACTTGCTTATTATCTCG
PtaNRX1.4       TTTTATGCTAACCTTGCATGTTGTGGTTGCATTACTTTCAATAACTTTCTTATCATCACG
PtaNRX1.1       TTCTCTGCTAACGTCGCATGTTGTTATTTCATTACTTACAATAACTTGCTTATTATCTCG
PtaNRX1.5       TTCTGTGCTAACATCGCATGTTGTTATTGCATTACTTTCAATAACTTGCTTATTATCTCG
PtaNRX1.2       TTCTGTGCTAACATCGCATGTTGTTATTGCATTACTTTCAATAACTTGCTTATTATCTCG
PtaNRX1.7       TTCTATGCTAACCTCGCATGTTGTTACCGCATTACTTTCAATAACTTTCTTATGATCACG
PtaNRX1.6       TTCTATGCTAACCTCGCATGTTGTTACCGCATTACTTTCAATAACTTTCTTATGATCACG
                ** * ******* * *********      ******** ********* ***** *** **

PtaNRX1.3       AATTGAATTCATTAGCAAAGG--------TATTTTTATCATTCAAATGGTTTACTC-TTG
PtaNRX1.4       AATTGAATTCATTAGCAAAGGG------TTTTTTTTATCATTTAAGTGGTTTGTTC-TTG
PtaNRX1.1       AATTGAATTCATTAGCAAAGG--------TATTTTTATCAGTCAAATGGTTTGCTC-TTG
PtaNRX1.5       AATTGAATTCATTAGCAAAGG--------TATTTTTATCATTCAAATGGTTTACTCTTTG
PtaNRX1.2       AATTGAATTCATTAGCAAAGG--------TATTTTTATCATTCAAATGGTTTACTCTTTG
PtaNRX1.7       AATTGAATTCATTAGCAAAGGGTTTTTTTTTTTTTTATCATTCAAATGGTGTGCTC-TTG
PtaNRX1.6       AATTGAATTCATTAGCAAAGGGTTTTTTTTTTTTTTATCATTCCAATGGTTTACTC-TTG
                *********************         * ********* *   * **** *   ** ***

PtaNRX1.3       ATTAGGTCCCAGTGTCTGATCTAGTTGGAAAGAACATTCTTCTTTACTTCTCAGCTCAAT
PtaNRX1.4       ATTAGGTCCGAGTGTCTGATCTAGTTGGAAAGAACATTCTTCTTTACTTCTCAGCTCAAT
PtaNRX1.1       ATTAGGTCCCAGTGTCTGATCTAGTTGGAAAGAACATTCTTCTTTACTTCTCAGCTCAAT
PtaNRX1.5       ATTAGGTCCCAGTGTCTGATCTAGTTGGAAAGAACATTCTTCTTTACTTCTCAGCTCAAT
PtaNRX1.2       ATTAGGTCCCAGTGTCTGATCTAGTTGGAAAGAACATTCTTCTTTACTTCTCAGCTCAAT
PtaNRX1.7       ATTAGGTCCCAGTGTCTGAACTAGTTGGAAAGAACATTCTTCTTTACTTCTCAGCTCAAT
PtaNRX1.6       ATTAGGTCCCAGTGTCTGAACTAGTTGGAAAGAACATTCTTCTTTACTTCTCAGCTCAAT
                ********* ********* ****************************************

PtaNRX1.3       GGTGCCCTCCTTGTCGTGCCTTTTTGCCCAAGCTAATTGAAGCATATCACACAATTAAAG
PtaNRX1.4       GGTGCCCTCCTTGTCGTGCCTTTTTGCCCAAGCTAATTGAAGCATATCACACAATTAAAG
PtaNRX1.1       GATGCCCTCCTTGTCGTGCCTTTTTGCCCAAGCTAATTGAAGCATATCACACAATTAAAG
PtaNRX1.5       GGTGCCCTCCTTGTCGTGCCTTTTTGCCCAAGCTAATTGAAGCATATCACACAATTAAAG
PtaNRX1.2       GGTGCCCTCCTTGTCGTGCCTTTTTGCCCAAGCTAATTGAAGCATACCACACAATTAAAA
PtaNRX1.7       GGTGCCCTCCATGTCGTGCCTTTTTACCCAAGCTAATTGAAGCATACCACACAATTAAAA
PtaNRX1.6       GGTGCCCTCCATGTCGTGCCTTTTTACCCAAGCTAATTGAAGCATACCACACAATTAAAA
                * ******** ************** ******************** ************

PtaNRX1.3       GAAAAGGCAATGCATTTGAGGTGATCTTCATCTCAAGTGACAGCGACCAATCCACCTTTG
PtaNRX1.4       GAAAAGGCAATGCATTTGAGGTGATCTTCATCTCAAGTGACAGCGATCAATCCACCTTTG
PtaNRX1.1       GAAAAGGCAATGCATTTGAGGTGATCTTCATCTCAAGTGACAGCGATCAATCCACCTTTG
PtaNRX1.5       GAAAAGGCAATGCATTTGAGGTGATCTTCATCTCAAGTGACAGCGATCAATCCACCTTTG
PtaNRX1.2       GAAAAGACAATGCATTTGAGGTGATCTTCATCTCAAGTGACAGAGATCAATCCACCTTTG
PtaNRX1.7       GAAAAGACAATGCATTTGAGGTGATCTTCATCTCAAGTGACAGAGATCAATCCACCTTTG
PtaNRX1.6       GAAAAGACAATGCATTTGAGGTGATCTTCATCTCAAGTGACAGAGATCAATCCACCTTTG
                ****** ************************************ ** *************

PtaNRX1.3       ACGAGTTCTATTCAGAAATGCCTTGGTTAGCCCTTCCATTTGGTGATGAAAGGAATCAAA
PtaNRX1.4       ACGAGTTCTATTCAGAAATGCCTTGGTTAGCCCTTCCATTTGGTGATGAAAGGAAACAAA
PtaNRX1.1       ACGAGTTCTATTCAGAAATGCCTTGGTTAGCCCTTCCATTTGGTGATGAAAGGAAACAAA
PtaNRX1.5       ACGAGTTCTATTCAGAAATGCCTTGGTTAGCCCTTCCATTTGGTGATGAAAGGAAACAAA
PtaNRX1.2       ACGAGTTCTATTCAGAAATGCCTTGGTTAGCCCTTCCATTTGGTGATGGAAGGAAACAAA
PtaNRX1.7       ACGAGTTCTATTCAGAAATGCCTTGGTTAGCCCTTCCATTTGGTGATGAAAGGAAACAAA
PtaNRX1.6       ACGAGTTCTATTCAGAAATGCCTTGGTTAGCCCTTCCATTTGGTGATGGAAGGAAACAAA
                ************************************************ ****** ****

PtaNRX1.3       TCCTGAGTCGGAAATTCAAAATTCAAGGCATTCCTGCAGCTGTATCGATTGGGCCAAGTG
PtaNRX1.4       TCCTGAGTCGGAAATTCAAAATTCAAGGCATTCCTGCAGCTGTATCGATTGGCCCAAGTG
PtaNRX1.1       TCCTGAGTCGGAAATTCAAAATTCAAGGCATTCTTGCAGCTGTAGCGATTGGCCCAAGTG
PtaNRX1.5       TCCTGAGTCGGAAATTCAAAATTCAAGGCATTCCTGCAGCTGTAGCGATTGGGCCAAGTG
PtaNRX1.2       TCCTGAGTCGGAAATTCAAAATTCAAGGCATTCCTGCAGCTGTAGCGATTGGCCCAAGTG
PtaNRX1.7       TCCTGAGTCGGAAATTCAAAATTCAAGGCATTCCTGCAGCTGTAGCGATTGGGCCAAGTG
PtaNRX1.6       TCCTGAGTCGGAAATTCAAAATTCAAGGCATTCCTGCAGCTGTAGCGATTGGGCCAAGTG
                ********************************* ********** ******* *******
```

```
PtaNRX1.3       GCCGGACCATTACGAAGGAAGCTCGGATGCACCTGACAGCTTACGGGGCAGATGCTTTTC
PtaNRX1.4       GCCGGACCATTACGAAGGAAGCTCGGATGCACCTGACAGCTTACGGGGCAGATGCTTTTC
PtaNRX1.1       GCCGGACCATTACGAAGGAAGCTCGGATGCACCTGACAGCTTACGGGGCAGATGCTTTTC
PtaNRX1.5       GCCGGACCATTACGAAGGAAGCTCGGATGCACCTGACAGCTTACGGGGCAGATGCTTTTC
PtaNRX1.2       GCCGGACCATTACCAAGGAAGCTCGGATGCACCTGACAGCTTACGGGGCAGATGCTTTTC
PtaNRX1.7       GCCGGACCATTACCAAGGAAGCTCGGATGCACCTGACAGCTTACGGGGCAGATGCTTTTC
PtaNRX1.6       GCCGGACCATTACCAAGGAAGCTCGGATGCACCTGACAGCTTAC-GGGCAGATGCTTTTC
                ************* ****************************** ***************

PtaNRX1.3       CATTTACCGAGGAGCATCTAAAGCAATTGGAGGAGGAGATTGAGGAAAAGGCAAAGGGGT
PtaNRX1.4       CATTTACCGAGGAACATCTAAAGCAATTGGAGGAGGAGATTGAGGAAAAGGCAAAGGGGT
PtaNRX1.1       CATTTACCGAGGAGCATCTAAAGCAATTGGAGGAGGAGATTGAGGAAAAGGCAAAGGGGT
PtaNRX1.5       CATTTACCGAGGAACATCTAAAGCAATTGGAGGAGGAGCTTGAGGAAAAGGCAAAGGGGT
PtaNRX1.2       CATTTACCGAGGAACATCTAAAGCAATTGGAGGAGGAGCTTGAGGAAAAGGCAAAGGGGT
PtaNRX1.7       CATTTACCGAGGAACATCTAAAGCAATTGGAGGAGGAGCTTGAGGAAAAGGCAAAGGGGT
PtaNRX1.6       CATTTACCGAGGAACATCTAAAGCAATTGGAGGAGGAGCTTGAGGAAAAGGCAAAGGGGT
                ************* ************************ *********************

PtaNRX1.3       GGCCAGAGAAAGTGAAACACGAACTT
PtaNRX1.4       GGCCAGAGAAAGTGAAACACGAACTT
PtaNRX1.1       GGCCAGAGAAAGTGAAACACGAACTT
PtaNRX1.5       GGCCAGAGAAAGTGAAACACGAACTT
PtaNRX1.2       GGCCAGAGAAAGTGAAACACGAACTT
PtaNRX1.7       GGCCAGAGAAAGTGAAACACGAACTT
PtaNRX1.6       GGCCAGAGAAAGTGAAACACGAACTT
                **************************
```

**Figure S2.** Multiple sequence alignments of *PtaNRX1* genes. Alignments were performed using MUSCLE (v3.8) provided by EMBL-EBI. Putative start and stop codons are shown in red and gRNA target sites in blue.
